# An open-source photogrammetry workflow for reconstructing 3D models

**DOI:** 10.1101/2023.03.12.532306

**Authors:** Chi Zhang, A. Murat Maga

**Affiliations:** Center for Development Biology and Regenerative Medicine, Seattle Children’s Research Institute, Seattle, Washington, United States of America; Division of Craniofacial Medicine, Department of Pediatrics, University of Washington, Seattle, Washington, United States of America

## Abstract

Acquiring accurate 3D biological models efficiently and economically is important for morphological data collection and analysis in organismal biology. In recent years, structure-from-motion (SFM) photogrammetry has become increasingly popular in biological research due to its flexibility and being relatively low cost. SFM photogrammetry registers 2D images for reconstructing camera positions as the basis for 3D modeling and texturing. However, most studies of organismal biology still rely on commercial software to reconstruct the 3D model from photographs, which impedes the adoption of this workflow in our field due the blocking issues such as cost and affordability. Also, prior investigations in photogrammetry did not sufficiently assess the geometric accuracy of the models reconstructed. Consequently, this study has two goals. First, we present an affordable and highly flexible structure-from-motion photogrammetry pipeline based on the open-source package OpenDroneMap (ODM) and its user interface WebODM. Second, we assess the geometric accuracy of the photogrammetric models acquired from the ODM pipeline by comparing them to the models acquired via microCT scanning, the de facto method to image skeleton. Our sample comprises fifteen *Aplodontia rufa* (mountain beaver) skulls. Using models derived from microCT scans of the samples as reference, our results show that the geometry of the models derived from ODM is sufficiently accurate for gross metric and morphometric analysis as the measurement errors are usually around or below 2%, and morphometric analysis captures consistent patterns of shape variations in both modalities. However, subtle but distinct differences between the photogrammetric and microCT-derived 3D models can affect the landmark placement, which in return affect the downstream shape analysis, especially when the variance within a sample is relatively small. At the minimum, we strongly advise not combining 3D models derived from these two modalities for geometric morphometric analysis. Our findings can be indictive of similar issues in other SFM photogrammetry tools since the underlying pipelines are similar. We recommend that users run a pilot test of geometric accuracy before using photogrammetric models for morphometric analysis. For the research community, we provide detailed guidance on using our pipeline for building 3D models from photographs.

## Introduction

Acquiring 3D biological models efficiently and economically can aid in data collection, research collaboration, and developing a comprehensive understanding of phenotypical variations and underlying biological mechanisms. Therefore, photogrammetry, the category of techniques that uses 2D photos for reconstructing 3D models with realistic texture, has become increasingly used in biological research (Giacomini et al., 2019; Waltenberger et al., 2021).

The conventional technique for the reconstruction of 3D models from photos is stereophotogrammetry, which typically involves multiple cameras and recording their positions (Duncan et al., 2022; Iglhaut et al., 2019; Westoby et al., 2012). Due to the non-invasive nature of photography, this technique has been used by clinicians in the past fifteen years to acquire craniofacial scans relatively quickly and safely from patients, as well as research in human craniofacial development and disorders (Al-Rudainy et al., 2018; Duncan et al., 2022; Heike et al., 2009, 2010; Weinberg et al., 2016).

Thanks to the fast advancement of computer vision techniques and infrastructure, structure-from-motion (SFM) photogrammetry tools have been rapidly developed and refined within the last decade (Fau et al., 2016; Giacomini et al., 2019; Morgan et al., 2019; Waltenberger et al., 2021). Unlike stereophotogrammetry, SFM photogrammetry estimates camera poses and positions from 2D image registration (Iglhaut et al., 2019; Westoby et al., 2012). Thus, a single camera can be used for data collection. Due to this flexibility, SFM photogrammetry has been widely used in 3D data acquisition for geographical and geological surveys, architectural preservation, and 3D modeling for archeological and paleontological sites (Wang et al., 2019; Westoby et al., 2012; Zimmer et al., 2018). In recent years, SFM photogrammetry has also been introduced to biological data collection for research, museum archives, and anatomical education as a flexible and low-cost tool (Giacomini et al., 2019; Knyaz et al., 2018; Lauria et al., 2022).

Most previous studies that evaluated SFM photogrammetry for biological research used commercial software, for which the costs might be substantial or yearly accruing (Fahlke & Autenrieth, 2016; Giacomini et al., 2019; Knyaz et al., 2018; Waltenberger et al., 2021).

Furthermore, although photogrammetry derived 3D models may look realistic due to the high-resolution texture, their geometric accuracy should be carefully assessed (Giacomini et al., 2019; Waltenberger et al., 2021). As pooling 3D data acquired by different digitization methods has been becoming more frequent, we urgently need to understand how the errors produced by these methods due to differences in the acquired geometry may disrupt detecting biological signals.

Previous tests of SFM photogrammetry have not incorporated sufficient evaluations of the geometric accuracy of the acquired textured models. Some studies only provided an overall evaluation and visualization of 3D model surface deviations between photogrammetric models and those acquired by other digitization methods (Buzi et al., 2018; Fahlke & Autenrieth, 2016; Fau et al., 2016; Waltenberger et al., 2021). Other studies that assess the performance of photogrammetric models in landmark-based morphometric analyses only showed that models acquired by digitization methods tended to cluster together in principal component plots (Buzi et al., 2018; Waltenberger et al., 2021). Only Giacomini et al. (2019) evaluated the error produced by digitization methods in a morphometric analysis using 3D models of bat skulls acquired by photogrammetry, CT scanning, and laser scanning. They suggested that, although photogrammetric models could be overall sufficiently accurate for multi-species evolutionary studies, researchers should be cautious in mixing data acquired by different digitization methods, especially when the sample showed limited variability.

Regarding measurement accuracy, Morgan et al. (2019) found that the average error between measurements taken on photogrammetric models of 45 human skulls and physical measurements fell below 2mm (around 2%). A 2-mm average measurement error is usually considered acceptable for anthropometric analysis (Katz & Friess, 2014; Morgan et al., 2019; Oriola et al., 2022; Stull et al., 2014). However, physical measurements may not be a good reference since they are more prone to operator errors compared to taking measurements on 3D models (Robinson & Terhune, 2017; Waltenberger et al., 2021; Weinberg et al., 2004). Other researchers focus on the consistency of measuring photogrammetric models compared to measuring physical specimens, laser or microCT scanned models (Jurda & Urbanová, 2016; Lauria et al., 2022; Lee & Gerdau-Radonic, 2020; Oriola et al., 2022; Weinberg et al., 2016).

In summary, our study has two main objectives. First, we presented a full workflow of photogrammetry from photography to 3D model post-processing using only free, open-source tools. The SFM photogrammetry was based on the WebODM, the convenient user interface of the open-source package OpenDroneMap (ODM) (Vacca, 2019, 2020; WebODM Authors, n.d.). We provided methods, tools, and detailed guidance to ease users’ introduction to photogrammetry. Second, we offered an assessment of the geometric accuracy of the ODM-derived models using both metric and landmark data. For this purpose, we used the models acquired from microCT scanning as the gold standard. We also focused on whether mixing photogrammetric and CT model can influence data analysis in a single-species sample with low variance (Giacomini et al., 2019).

## Materials and Methods

### Materials

Our sample comprised of fifteen adult mountain beaver (*Aplodontia rufa*) skulls. Fourteen skulls were provided by the courtesy of Burke Museum of Natural History, Seattle, Washington, USA. An additional skull was provided from the personal collection of one of the authors (AMM). Burke Museum accession numbers can be found in Table S1 in the supplemental material.

### 3D model reconstruction using OpenDroneMap and WebODM

We used a low-end DSLR camera mounted on a tripod, a remote-controllable turntable, and a lightbox for photography. The specifics of the photography setup and imaging protocol, along with the open-source software to control the data acquisition can be found in the workflow guidance in the workflow instructions of the supplementary material.

SFM photogrammetry was performed using WebODM, the graphic user interface of the open-source package ODM (Vacca, 2020; WebODM Authors, n.d.). The essential steps of SFM photogrammetry in ODM can be summarized as:

1. Image registration based on feature matching.
2. Multi-view stereo: this process reconstructs camera poses and positions based on image registration and the camera metadata. The output is a sparse point cloud.
3. Dense point cloud creation: using the information and the sparse point cloud from the last step to generate a dense point cloud.
4. Modeling and texturing: by default, ODM uses Poisson reconstruction for 3D modeling based on the dense point cloud and then maps texture to the model.
5. Model scaling: In ODM, it is possible to scale 3D models to their physical sizes using Aruco Markers.

### 3D Model reconstruction using microCT scanning

We used a Bruker/Skyscan 1076C microtomography microCT scanner to acquire 3D scans of mountain beaver skulls at 35-micron resolution. These scans were converted into 3D models using Segment Editor module of 3D Slicer (Kikinis et al., 2014) and SlicerMorph (Rolfe et al., 2021) as microCT imaging is considered the “gold standard” to acquire geometrically accurate models. For detailed microCT scanning protocols, please see Table S2 in the supplementary material.

### Landmark collection

The photogrammetric models (ODM-derived hereafter) and micro-CT scanned models (CT-derived hereafter) were imported into 3D Slicer for landmark collection (Kikinis et al., 2014). The same operator (CZ) annotated 29 landmarks on each model (Figure S1, Table S3). The procedure was replicated three times. The average of the three replicates was the final landmark set for metric and morphometric analysis, which we refer as mean landmark dataset in sections below. Replicates were also used to calculate intraobserver error for each method. All error calculation and analysis were performed using statistical language R (R Core Team, 2017).

### Measurement errors and accuracy

Because the ODM-derived models had been automatedly scaled using the Aruco markers to approximate real-life sizes, the Euclidean distances between landmarks were directly used to represent the actual measurements. We calculated seventeen linear measurements for each model based on the mean landmark set: six anteroposterior length measurements, nine bilateral width measurements between pairwise landmarks, and two height measurements (Table S4). We reported inter-method measurement error by subtracting the ODM-derived measurement from the CT derived one, and taking its absolute value: error_inter-method_ = (abs(CT_measure_ – ODM_measure_)). The inter-method error for each measurement was also converted to the percentages of corresponding CT-derived measurement (inter-method percent error in the following text): error_inter-method_/CT_measure_ × 100.

The accuracy of the ODM-derived measurements was determined as how similar they were to the gold standard, the CT-derived measurements. For this purpose, we conducted two-sided t-test to assess whether the mean CT and ODM-derived values for each of the 17 measurements were significantly different. If the mean CT and ODM-derived measurements were not significantly different (p > 0.05) for the majority of the 17 measurements, the accuracy of the ODM-derived measurements could be considered as acceptable for statistical analysis.

### Errors in geometric morphometric analysis

We first assessed the errors created by the two digitization methods, photogrammetry (ODM) and CT scanning, in a morphometric analysis based on a mixed dataset with landmarks annotated on the models derived from these two methods. To do this, we performed a Procrustes ANOVA with random residual permutation (RRPP) based on a joint Generalized Procrustes Analysis (GPA) of all three ODM and CT-derived landmark sets using the geomorph R package (Adams et al., 2021; Collyer et al., 2015). GPA registers landmark configurations by removing size, location, and orientation factors (Zelditch et al., 2012). Similar to ordinary ANOVA, Procrustes ANOVA is designed to quantify variances explained by different factors within a landmark dataset registered by GPA and test whether these variances are significant (Goodall, 1991; Klingenberg & McIntyre, 1998).

Our linear model (Procrustes coordinates ∼ ID + Method + Replicates + ID:Method) contained three variables and an interaction term: 1) ID: individual variations based on labeling the ODM and CT-derived models with the same ID, 2) Method: the mean error (or systematic error) caused by the two digitization methods: photogrammetry and CT scanning, 3) Replicates: intraobserver errors from three landmark trials, and 4) ID by Method: the interaction between digitization methods and ID. This fourth term, the interaction, represents the variance explained by the variation in the errors caused by the two digitization methods across individuals. In other words, this interactive factor quantifies the random errors associated with digitization methods occurred at the individual level. R square of each factor quantifies the proportion of variance it explains. If the p-value of each factor is smaller than 0.05, the factor explains a significant amount of variance. In general, if the landmarks are carefully annotated and the digitization methods yield highly consistent landmark sets, the “ID” factor should account for nearly all the total variance with an extremely small p-value.

Additionally, separate GPAs of the ODM and CT-derived mean landmark datasets were performed to assess whether they captured similar patterns of shape variation and resulted in similar morphospace. First, we computed the correlation coefficients between all pairs of Procrustes distances of the ODM and CT-derived datasets. The Procrustes distance is the root of the sum of squares between two landmark configurations, thus representing their overall shape difference. Second, we calculated the correlation coefficient between the principal component (PC) scores of the first five PCs. Principal Component Analysis (PCA) is commonly used to reduce the dimensionality of the hyper-dimensional morphospace by generating principal components (PCs) ordered by the variances they explained. Usually, the first few PCs are used to summarize the patterns and magnitude of the overall variations within a sample. Thus, the correlation between corresponding PCs assesses the similarity between the morphospace of the ODM and CT datasets. Overall, if the correlation coefficient (r) exceeded 0.8 and the p-value was below 0.05, the two variables were considered as strongly correlated. Ideally, if the morphospaces yielded from the CT and ODM-derived landmark sets are highly consistent, the correlation coefficients of Procrustes distances and first five PCs should all be close to 1.

## Results

### Quality of the ODM-derived textured models

The raw photos and the output ODM-derived models (OBJ file) are available publicly in an online repository (https://osf.io/b39yx/, DOI: 10.17605/OSF.IO/B39YX) (Figure S1). Overall, the quality of the textured models was high and sufficient for visual comparison and landmark annotation. The ODM-derived textured models have around 400,000 to 700,000 vertices. There were some black polygons (noise) attached to thin edges and structures, such as the external and internal surfaces of the zygomatic arches and the anterior margin of the nasal bones (Figure S3). However, they, in general, did not influence landmarking (Figure S1). We only used functions for removing selected polygons and isolated polygons in MeshLab to delete the black polygons attached at the external surface of the zygomatic arches because they may influence landmark annotation in a few specimens (see section 6.1 in the workflow instructions of the supplementary material).

The ODM-derived models were essentially watertight (Figure S2). The holes and foramina, such as incisive foramina and even foramen magnum, in the ODM-derived models were closed. Furthermore, the sutures were also marked by texture and did not show on the surface of the meshes. The fissures at the two sides of the occipital bones were also fused. These structures were delineated only by texture. Thus, we relied on using texture to place the landmarks on these structures.

### Processing time for ODM-derived models

When syncing the turntable with the camera, it took approximately two minutes to photograph each of the six sets of photos (32 to 64 photos). Overall, taking photos for one specimen took around 20 to 30 minutes, which included setting up specimens into different orientations and adjusting the camera focus ring. After training with three to four specimens, the time for taking photos of one specimen could drop to around 20 minutes. It took less than 10 minutes per specimen for photo preprocessing, such as using a custom script in 3D Slicer to create a rectangular box for an initial background masking for each set of photos (section 4 in the workflow instructions of the supplementary material). Using a cloud server based ODM, creating a 3D model from photographs took about two to three hours. We configured the WebODM to allow running two tasks concurrently to increase the throughput. Overall, the fifteen beaver models took approximately twenty hours to process.

### Metric errors and accuracy

The overall mean inter-method measurement error was 0.550mm. The mean inter-method errors of the seventeen measurements ranged from 0.138mm to 1.376 mm (Figure 2 and Table S6). When converting the inter-method errors to the percentages of the corresponding CT-derived measurements, the overall mean inter-method percent error was 1.760%. The mean inter-method percent errors of the seventeen measurements ranged from 0.998% to 3.057% (Figure 2, Table S6 and Table S7). 95% of the inter-method percent errors fell below 4%. The large inter-method precent errors were primarily due to the measurements are small scaled.

**Figure 2.**
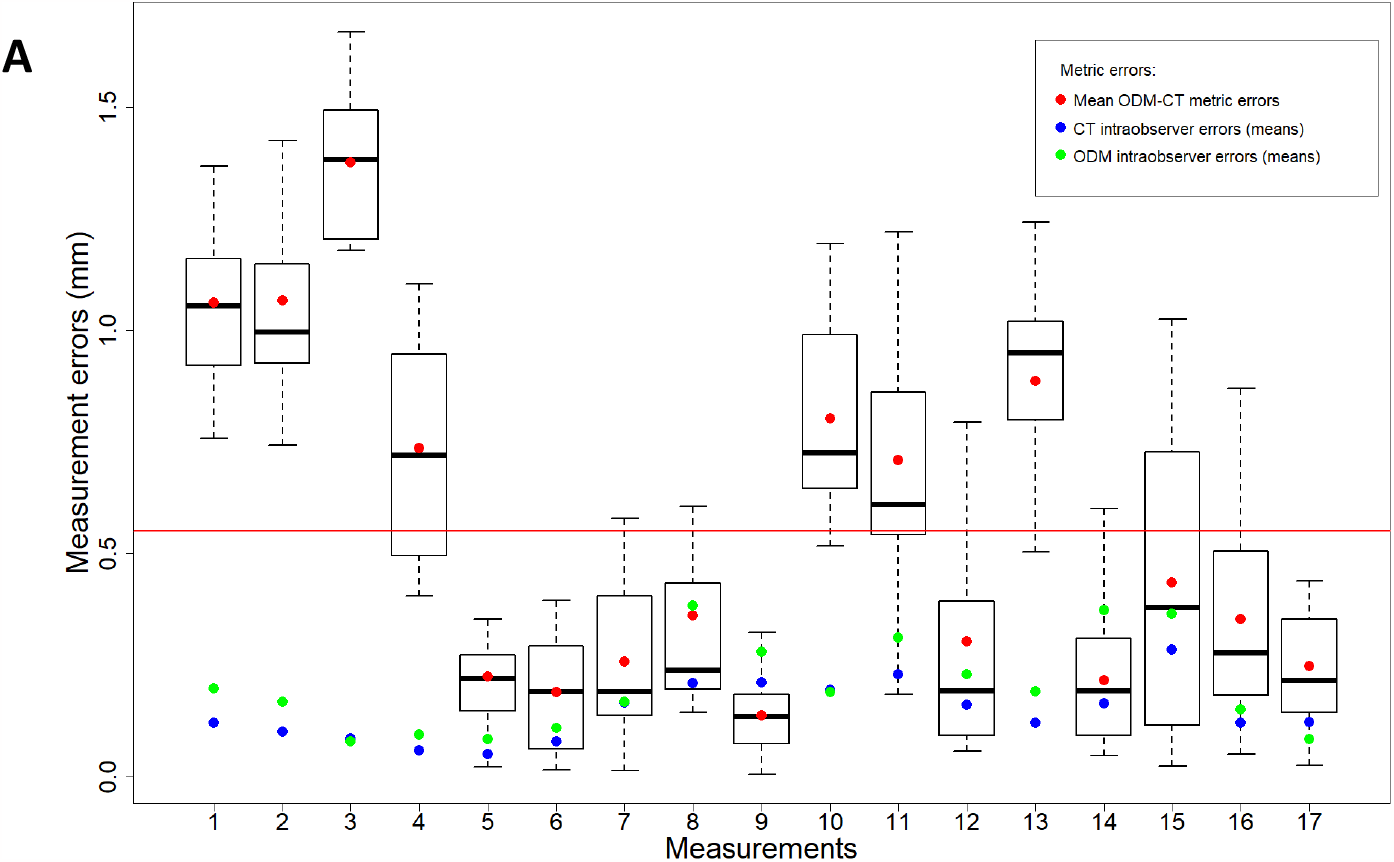
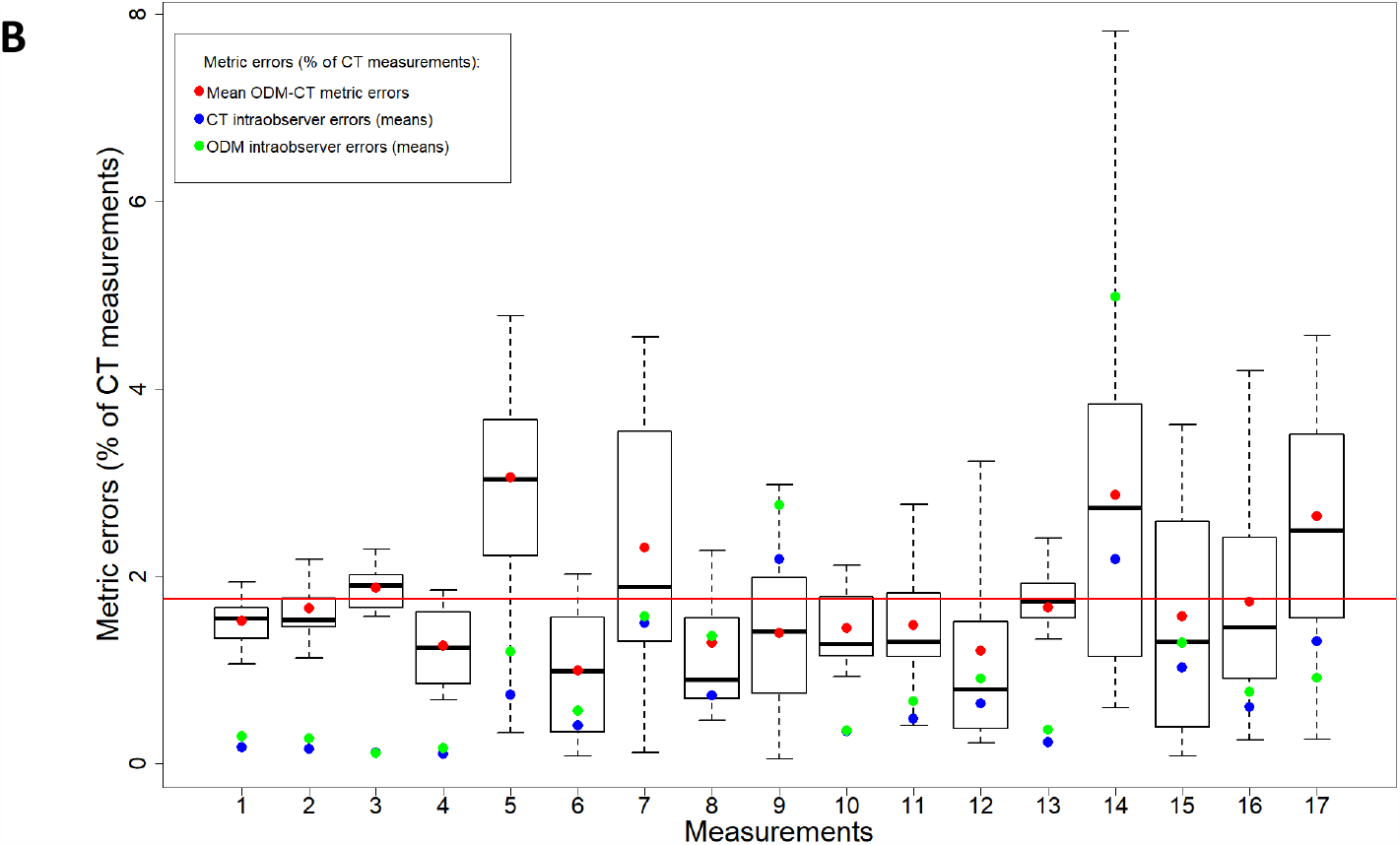
Inter-method measurement errors based on the automatedly scaled ODM-derived datasets. (A) Inter-method measurement errors in absolute values. (B) Inter-method measurement errors as percentages of the corresponding CT measurements. Each column represents the inter-method errors between the CT and ODM-derived sets for each of the 17 measurements. Red dots: mean inter-method measurement errors. Blue dots: mean CT-derived intraobserver errors. Green dots: mean ODM-derived intraobserver errors. Red horizontal line: overall mean ODM-CT measurement error. For both modalities, intraobserver measurement errors were calculated by averaging the errors between measurements derived from the mean landmark set and each replicate. The CT and ODM-derived intraobserver errors are not significant different for any measurement based on two-sided Welch t-tests (p > 0.05).

For example, the largest inter-method percent error (7.824%) was from the specimen 82409’s measurement 14, a small-scaled measurement between two premolars. However, the corresponding absolute inter-method measurement error was not exceptionally large (0.601mm). Two-sided Welch t-tests showed that the mean CT and ODM-derived values were not significantly different for all measurements except for measurement 3 (p > 0.05). Thus, the accuracy of the ODM-derived measurements was acceptable for statistical analysis.

### Geometric Morphometric analysis

Procrustes ANOVA based on a joint GPA of the three CT and ODM-derived landmark replicates showed that the individual variations (the factor ‘ID’) accounted for 90.8% of the total variance (R-square) (Table 1). Though statistically significant, intraobserver errors (the factor ‘replica’) from three rounds of manual landmarking only accounted for 0.23% of the total variance.

**Table 1.** Procrustes ANOAVA after a joint GPA of the ODM and CT datasets.

|  | Degree of Freedom | Sum of Square | Mean square | R square | F value | P value |
| --- | --- | --- | --- | --- | --- | --- |
| ID | 14 | 0.107 | 0.00761 | 0.908 | 91.049 | $1 \times 10^{-4}$ |
| Method | 1 | 0.00127 | 0.00127 | 0.0108 | 15.171 | $1 \times 10^{-4}$ |
| Replica | 2 | 0.000270 | 0.000135 | 0.00230 | 1.615 | 0.0485 |
| ID: method | 14 | 0.00443 | 0.000317 | 0.0378 | 3.786 | $1 \times 10^{-4}$ |
| Residual | 58 | 0.00485 | 0.0000836 | 0.0413 |  |  |
| Total | 89 | 0.117 |  |  |  |  |

Therefore, the intraobserver errors can be considered as minimal. The factor ‘method’, the mean difference between the CT and ODM-derived landmark datasets, accounted for a small but significant amount of the total variance (1.08%). The interaction between individual specimens and the digitization methods, as represented by ‘ID : method’, explained 3.78% of the total variance and was highly significant. This showed that the variation in the difference between two digitization methods across individuals accounted for a small but highly significant amount of the total variance. In other words, digitization methods also created significant random errors at the individual level in addition to systematic error.

The correlation analysis of pairwise Procrustes distances and PC scores from the separate GPAs of the CT and ODM-derived landmark sets confirmed deviations caused by the two digitization methods in morphometric analysis. Though strongly correlated (p-value < 0.001 and correlation coefficient r > 0.8), the correlation coefficient between the CT and ODM-derived pairwise distances fell below 0.9. (Figure 3A). None of the correlation coefficients between the scores of the first five CT and ODM-derived PCs, which explained 70.7% and 72.7% of the total variance, respectively, exceeded 0.9 (Figure 3B). In particular, the correlation coefficient between the CT and ODM-derived PC1 scores fell below 0.8.

**Figure 3.**
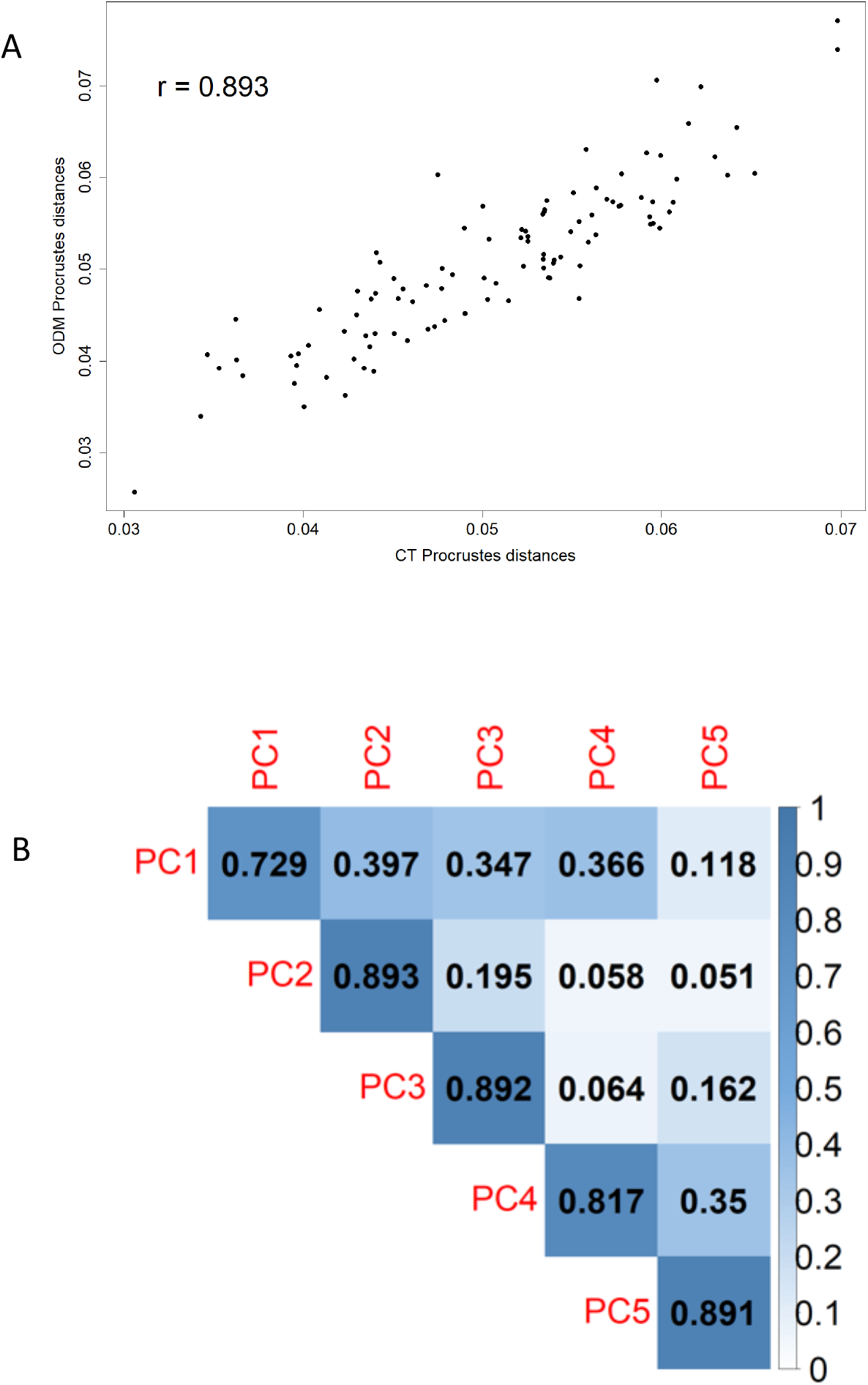
Correlations of morphometric variables from separate GPAs of the ODM and CT datasets. (A) Correlations of pairwise Procrustes distances. (B) Correlations of the scores of the first five PCs.

## Discussion

### Geometric accuracy of the ODM-derived models

Our study showed the linear measurements taken on the ODM-derived models were in general consistent with the CT-derived ones. Therefore, the ODM-derived models were sufficiently accurate for statistical analyses using linear measurements. The mean inter-method measurement error fell below 2%. Morgan et al. (2019) considered that an average error of 2% percent was acceptable for human osteometric analysis. However, in our study, the inter-method measurement errors still exceeded the intraobserver errors. If the ODM and CT-derived models were nearly identical, the inter-method errors should be comparable with intraobserver error. Therefore, users should still be cautious about analyzing subtle variations, such as individual variations, using measurements taken on the ODM-derived models.

The performance of ODM-derived landmark sets in morphometric analysis yielded mixed results. In general, we concur with the previous studies (e.g., Giacomini et al., 2019; Katz & Friess, 2014; Waltenberger et al., 2021) that Landmarks collected from the ODM-derived (and other photogrammetry-derived) models can be sufficient for capturing overall trends within a sample. In other words, CT and ODM-derived models were overall highly consistent in geometry. However, the landmarks placed on the ODM-derived (and other photogrammetry-derived) 3D models may not be ideal for investigating subtle variation, such as variation within a species with limited variability. Nearly 5% of variance was associated with digitization methods in a mixed CT and ODM-derived landmark set. The CT and ODM-derived landmark sets also showed clear deviations in morphospace. Consequently, the differences in the model geometry created by two digitization methods may conceal subtle but biologically meaningful signals. Moreover, in addition to the subtle but significant systematic error created by the CT and ODM methods, the two digitization methods generated significant random error across individuals. In other words, landmarks placed on CT and ODM-derived models can produce different levels of error, subject to the condition of each individual specimen. The random errors created by digitization errors are usually hard to predict and control, thus can further complicate the results of morphometric analysis based on landmarks placed on ODM-derived models. Therefore, we concur with Giacomini et al. (2019) and advice not to mix landmark sets taken from the CT and ODM-derived models for morphometric analysis, in particular when the focus of the analysis is individual variation.

It should be noted that the results of error testing in our study (as well as in any study of error based on landmark data) depended on the landmark choice, sample, and experience of the operator (Giacomini et al., 2019). For example, our study was based on a single species with limited variability. The impact of digitization methods on morphometric analysis may be reduced if our sample contains well differentiated groups, such as different species an evolutionary study (Giacomini et al., 2019). Nevertheless, our findings can still indicate similar issues in other SFM photogrammetry tools since the underlying pipelines are similar. Therefore, we recommend users to run a pilot study to determine whether models acquired by photogrammetry are appropriate for metric and morphometric analyses.

### The importance of photography settings for photogrammetry

Our photogrammetry pipeline completely relies on open-source software, thus further bringing the cost down compared to previous studies using commercial software. The downside of any photogrammetry pipeline is that it can be laborious. The photo taking and preprocessing can be highly time-consuming and is subject to the experience of users (Fahlke & Autenrieth, 2016; Fau et al., 2016; Waltenberger et al., 2021). We offered a variety of tools to ease the process of photography and improve the repeatability of data collection, such as syncing the camera with the turntable and using Depth-of-Field to set up camera parameters (see supplementary material). With sufficient training, the time it took to take 320 pictures for each specimen can be reduced to around 20 minutes.

Here we present several tips and methodological concerns for taking photos (also see the workflow instructions of the supplementary information). The first is about the sharpness of the photos. For camera parameters, the F stop values determine the sharpness of pictures. We set up the F stop values between F13 to F16. However, based on our earlier experiments, F/20 to F/22 could still achieve sharp enough photos for high-quality texturing based on our relatively low-end DSLR camera.

It is critical to cover sufficient detail of the specimen by taking pictures from different angles. We used six sets of 320 photos for each specimen. We first experimented with taking three sets of vertically oriented specimens and found that this setting is better for capturing the trunk of the skull. We then added two more sets of photos of a horizontally placed specimen to better capture the lateral structures, such as zygomatic arches and the external acoustic meatus.

Users can experiment with reconstructing models with photos taken from different angles to see which angles do better in capturing certain aspects of the objects. We recommend taking more than 32 pictures (i.e., no more than 11.25 degrees difference per rotation) for each full circle of turntable rotation to make sure sufficient overlapping between adjacent pictures exists.

Ambient light should be provided to expose sufficient details of the specimen. However, eliminating shadows is not possible. Shadow is not an issue if covered structures are sufficiently exposed in other photos. Glare and reflective surfaces would also cause problems for photogrammetry. Therefore, overly intense lightning should be avoided. In general, our beaver skulls were smooth and reflective in a few areas, but we did not encounter problems with model reconstruction.

### Photogrammetric process and other methodological concerns

WebODM is a convenient interface to run the ODM package for photogrammetric reconstruction (Vacca, 2020; WebODM Authors, n.d.). Users only need to submit photos and specify parameters to run the process. We provided a reference parameter setting in the pipeline guidance (see section 5.2 in the workflow instructions of the supplementary material).

Reducing background noise is critical to ensure successful photogrammetric reconstruction and high-quality textured modelling (Matuzevičius & Serackis, 2021). Simply covering the background with the same color is helpful but insufficient. For example, the thin structures, such as zygomatic arches, might be covered by black polygons that may obscure placing landmarks (Figure S4). The same difficulty in reconstructing thin structures has also been reported by other researchers using commercial software (Giacomini et al., 2019). ODM has a deep learning background removal algorithm to mask the background automatedly. This option must be checked to ensure consistent successful reconstruction for our photogrammetric reconstruction and substantially improve the quality of the textured model, especially the thin structures (Figure S4) (see section 5.2 in the workflow instructions of the supplementary material). Some pre-processing of acquired photos was still necessary. To prepare for the background removal, we used a script to call functions in 3D Slicer to create a bounding box that contains the specimen (see section 4.1 in the workflow instructions of the supplementary material). We then masked the area outside the box as black. This step of preprocessing could reduce the burden of background removal during the model reconstruction process in ODM and ensure consistent success for each sample. This step was also necessary to remove the putty that fixed the specimen vertically, or else the putty would be reconstructed as well. Note that the small areas of the specimen can be masked if the same areas are provided in other pictures.

Finally, we recommend using Aruco markers not just for automated scaling but also for ensuring successful textured model reconstruction in ODM (see section 2.3 and 3 in the workflow instructions of the supplementary material). Based on our experience, the model reconstruction task may fail due to the inability to estimate a scale from the input photos.

## Conclusion

We presented an open-source SFM photogrammetry pipeline using WebODM to acquire textured models from biological specimens. We offered a variety of tools and detailed guidance to simplify the data collection process and ensure consistency and repeatability in model reconstruction. Comparing to the “gold standard” microCT-derived models, the reconstructed textured models were sufficiently accurate for assessing overall shape differences and conducting metric analysis. However, differences between the CT and ODM-derived models, though small, could still significantly impact detailed morphological analysis. While our analysis is specific to the open-source ODM photogrammetry toolkit, given the SFM photogrammetry pipelines are similar across different programs, our finding is not unique to ODM, but likely indicative of similar issues in other photogrammetry tools. Users should be cautious about mixing photogrammetric models with models acquired by other digitization methods.

## Supporting information

Supplementary tables, figures, and workflow instructions

## Data availability

CT and ODM-derived models (texture mapped to the model as vertex color) in PLY format, sample photo set for one mountain beaver skull, and landmark datasets used in the analysis are available at https://osf.io/b39yx/ (DOI: 10.17605/OSF.IO/B39YX).

## Acknowledgement

We thank Jeff Bradley (Burke Museum) for arranging the loan of mountain beaver skulls used in this research. We thank Stephen Mather for his help on optimizing the workflow for ODM and help us configure WebODM on the cloud. We also acknowledge the communities of both OpenDroneMap and 3D Slicer for their support and contribution to our research. Parts of this research was funded by National Science Foundation [OAC 2118240] (Imageomics Institute) and ACCESS [BIO180006] (SlicerMorphCloud) to AMM.

