## Supplementary tables, figures, and workflow instructions for "An open-source photogrammetry workflow for reconstructing 3D models": supplement_tables_figures.pdf

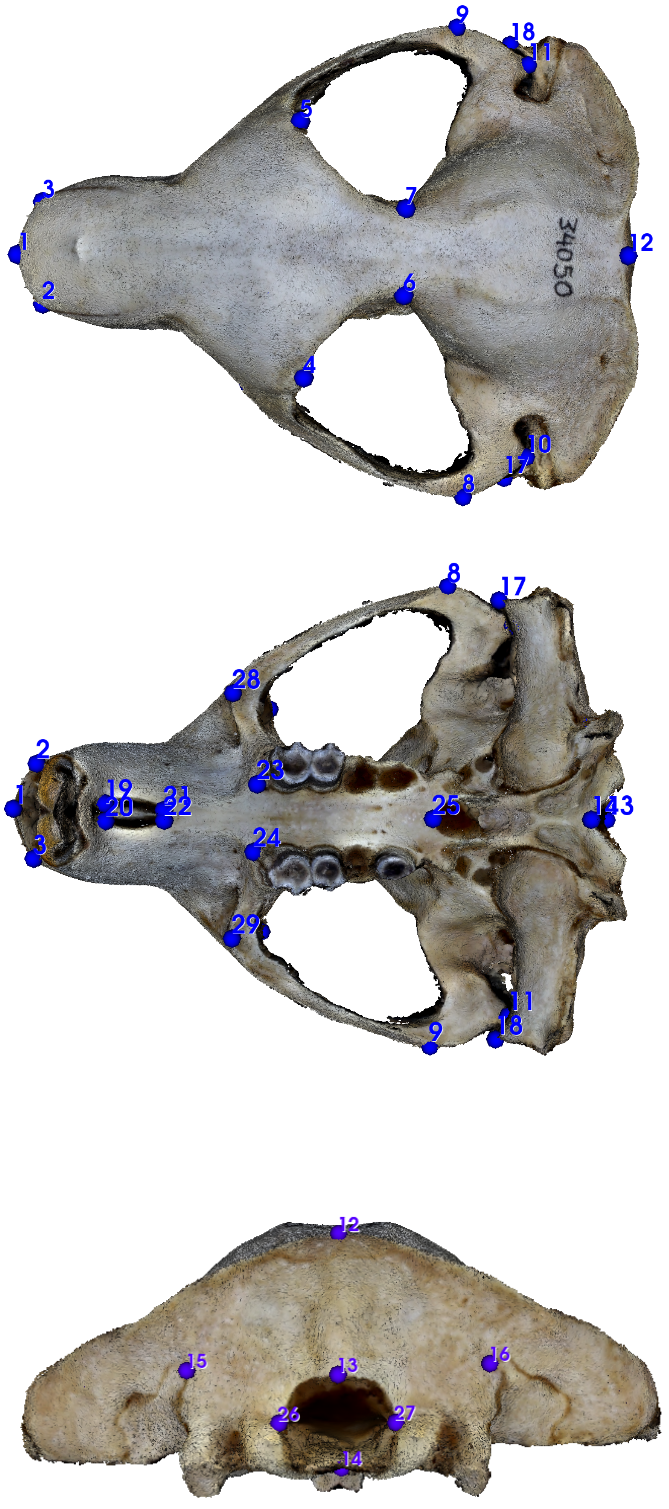


**Figure S1. Twenty-nine Landmarks on an ODM-derived textured model.** Visualized in 3D Slicer.

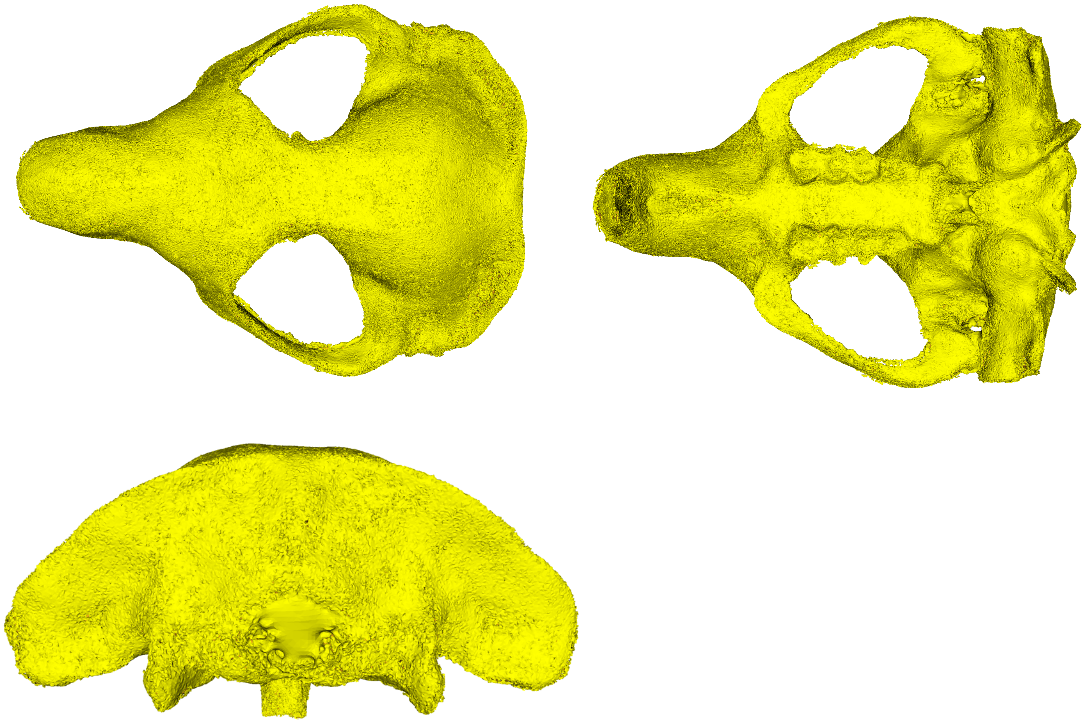

**Figure S2. ODM-derived models (OBJ) without texture.** Visualized in 3D Slicer.

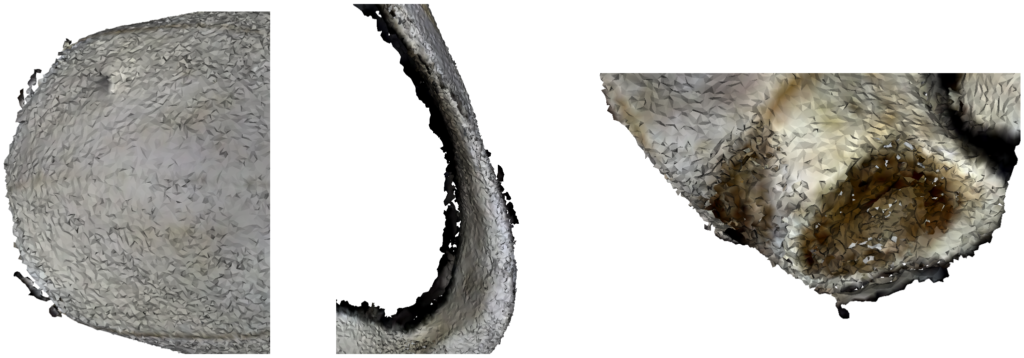

**Figure S3. Black polygons (noise) at the surface of thin structures in some specimens, such as the anterior nasal margin (left), inner and outer surface of the zygomatic arch (middle), and the margin of the external ear tube (right).** Visualized in 3D Slicer.

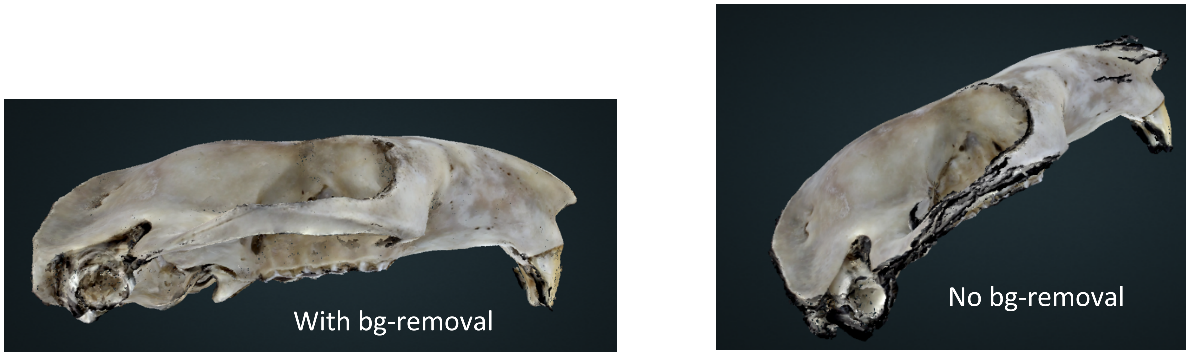


**Figure S4. The ODM-derived models of the same specimen (82710) using the background removal algorithm ("bg-removal") and without using "bg-removal".** Visualized in WebODM.

**Table S1. Fourteen specimens acquired from Burke Museum of Natural History, Seattle, WA, USA.**

31870, 34050, 34072, 34073, 34081, 34083, 35998, 39384, 39385, 39867, 39867, 79558, 79564, 82409, 82710

**Table S2. Protocols for using the Skyscan 1076C microtomograph (Bruker) microCT scanner for the mountain beaver sample.**

| | |
|---|---|
| Flat field setting | Aluminum filter, 0.55mm, acquire single, uncheck camera central position |
| Resolution | 35 micron |
| Scanning mode | Wide |
| NReconn density histogram range | 0.004 to 0.07 |

**Table S3. Landmark definitions.**

| Landmarks | Definition |
|---|---|
| 1 | Nasal: intersection of nasal bones, rostral point (placed at the left nasal bone) |
| 2, 3 | Anterior-most point at intersection of premaxillae and nasal bones (placed at the maxilla), left and right |
| 4, 5 | Intersection of frontal process of maxilla with frontal and lacrimal bones (the corner at the lateral side of the notch), left and right |
| 6, 7 | Most medial point of the temporal line, left and right |
| 8, 9 | Most lateral point on the zygomatic arch, left and right |
| 10, 11 | Most posterior point at the zygomatic arch, left and right |
| 12 | The midsagittal point at the superior margin of the nuchal plane |
| 13 | Opisthion: midsagittal point on the superoposterior margin of the foramen magnum |
| 14 | Basion: midsagittal point anteroinferior margin of the foramen magnum |
| 15, 16 | Medial end of the fissure at the nuchal plane, placed on the nuchal plane, left and right |
| 17, 18 | Most anterior point at the external margin of the acoustic meatus, left right |
| 19 | Most anterior point of the anterior palatine foramen, left and right |
| 21, 22 | Most posterior point of the posterior palatine foramen, left and right side |
| 23, 24 | Most anterior point at the base of the reduced first premolar, left and right |
| 25 | Posterior nasal spine at the midline |
| 26, 27 | Intersection of the right occipital condyle and the foramen magnum, left and right |
| 28, 29 | Anterior most point at the articular eminence, left and right |

**Table S4. Measurements.**

| | |
|---|---|
| **Length** | |
| Measurement 1 | Landmark 1-12 |
| Measurement 2 | Landmark 1-11 |
| Measurement 3 | Landmark 1-13 |
| Measurement 4 | Landmark 14-19 |
| Measurement 5 | Landmark 19-21 |
| Measurement 6 | Landmark 14-25 |
| **Width** | |
| Measurement 7 | Landmark 2-3 |
| Measurement 8 | Landmark 4-5 |
| Measurement 9 | Landmark 6-7 |
| Measurement 10 | Landmark -9 |
| Measurement 11 | Landmark 10-11 |
| Measurement 12 | Landmark 15-16 |
| Measurement 13 | Landmark 17-18 |
| Measurement 14 | Landmark 23-24 |
| Measurement 15 | Landmark 28-29 |
| **Height** | |
| Measurement 16 | Landmark 12-14 |
| Measurement 17 | Landmark 13-14 |

**Table S6. The mean inter-method and intraobserver errors for each of the 17 measurements.**

| | Inter-method error | | CT-derived intraobserver error | | ODM-derived intraobserver error | |
|---|---|---|---|---|---|---|
| Measurements | Absolute (mm) | Percent (%) | Absolute (mm) | Percent (%) | Absolute (mm) | Percent (%) |
| 1 | 1.061 | 1.525 | 0.120 | 0.173 | 0.197 | 0.288 |
| 2 | 1.067 | 1.652 | 0.0994 | 0.155 | 0.167 | 0.264 |
| 3 | 1.376 | 1.875 | 0.0854 | 0.116 | 0.0782 | 0.108 |
| 4 | 0.735 | 1.256 | 0.0583 | 0.0970 | 0.0941 | 0.163 |
| 5 | 0.224 | 3.057 | 0.0589 | 0.732 | 0.0842 | 1.197 |
| 6 | 0.189 | 0.988 | 0.0780 | 0.402 | 0.108 | 0.564 |
| 7 | 0.258 | 2.301 | 0.165 | 1.503 | 0.167 | 1.567 |
| 8 | 0.361 | 1.290 | 0.208 | 0.729 | 0.382 | 1.355 |
| 9 | 0.137 | 1.391 | 0.211 | 2.180 | 0.279 | 2.757 |
| 10 | 0.803 | 1.448 | 0.194 | 0.346 | 0.189 | 0.347 |
| 11 | 0.710 | 1.478 | 0.228 | 0.479 | 0.310 | 0.660 |
| 12 | 0.302 | 1.199 | 0.160 | 0.637 | 0.228 | 0.909 |
| 13 | 0.886 | 1.660 | 0.121 | 0.227 | 0.190 | 0.362 |
| 14 | 0.215 | 2.865 | 0.164 | 2.183 | 0.373 | 4.987 |
| 15 | 0.435 | 1.567 | 0.284 | 1.021 | 0.363 | 1.287 |
| 16 | 0.352 | 1.723 | 0.120 | 0.602 | 0.151 | 0.767 |
| 17 | 0.247 | 2.640 | 0.122 | 1.307 | 0.0836 | 0.917 |

**Table S7 Quantiles of the inter-method measurement errors.**

| Quantiles | 0 | 25% | 50% | 75% | 100% |
|---|---|---|---|---|---|
| Absolute errors (mm) | 0.00540 | 0.185 | 0.422 | 0.909 | 1.773 |
| Percent error (%) | 0.0486 | 1.056 | 1.553 | 2.068 | 7.824 |

**Table S8. P-values of the two-sided Welch t-tests between mean CT and ODM-derived measurements.**

| Measurements | P-value |
|---|---|
| 1 | 0.0761 |
| 2 | 0.0765 |
| 3 | 0.0207 |
| 4 | 0.175 |
| 5 | 0.341 |
| 6 | 0.712 |
| 7 | 0.510 |
| 8 | 0.555 |
| 9 | 0.875 |
| 10 | 0.326 |
| 11 | 0.170 |
| 12 | 0.832 |
| 13 | 0.173 |
| 14 | 0.991 |
| 15 | 0.884 |
| 16 | 0.233 |
| 17 | 0.200 |
