## Supplementary tables, figures, and workflow instructions for "An open-source photogrammetry workflow for reconstructing 3D models": supplement_workflow instructions.pdf

### Instructions for photogrammetry pipeline and data collection

#### 1. Prerequisites: Setting up the WebODM, finding help and additional software.

There are multiple ways ODM and WebODM can be installed. We opted to use the docker-based installation of the WebODM on our cloud server as it is the simplest to start. Please follow the instruction on setting up the WebODM at: https://github.com/OpenDroneMap/WebODM/#getting-started

The official documentation and tutorials of ODM (and WebODM) can be found at: https://docs.opendronemap.org/.

The user community form of ODM is at: https://community.opendronemap.org/

The postprocessing of the textured model and landmark data collection requires MeshLab and 3D Slicer with the SlicerMorph extension. MeshLab can be downloaded from https://www.meshlab.net/. For installing 3D Slicer and SlicerMorph, please see https://github.com/SlicerMorph/SlicerMorph#installation

A python programming environment is necessary if the user wants to scale the specimen using Aruco markers. See description in Section 3.

**Equipment used:**

- A DSLR camera (Canon EOS Rebel T6)
- A programmable turntable that can be controlled by a remote controller to set up how many steps it takes to finish a full circle and can be synced with a camera (The turntable we used was ComXim 12.6inch turntable with a shutter cable)
- Portable photo studio box (Amazon Basics) and lighting system (two UBeeSize V107 lights)
- Tripod (Amazon Basics)
- Putty
- A vertical stand for fixing the putty
- A remote shutter cable to connect the turntable with the camera, and a USB cable to connect the camera to the computers

#### 2. Photography:

##### 2.1 Equipment setup

Setup the turntable and lighting in the photo studio. Attach the camera to a tripod that allows three degrees of freedom for movement.

- We advise coverings the side walls and the studio floor in black and also paint or cover the surface of turntable black to minimize reflection and glare.
- Connect the camera and the turntable via the appropriate shutter cable.

- The camera is also synced with the computer using the DigiCamControl open-source software that works with Nikon and Canon cameras (http://digicamcontrol.com/). This allows direct transfer of photos to the computer from camera.

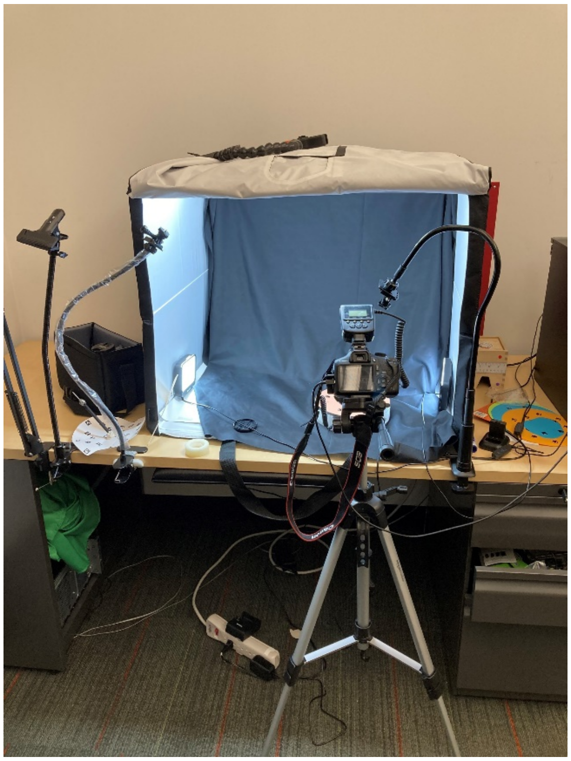

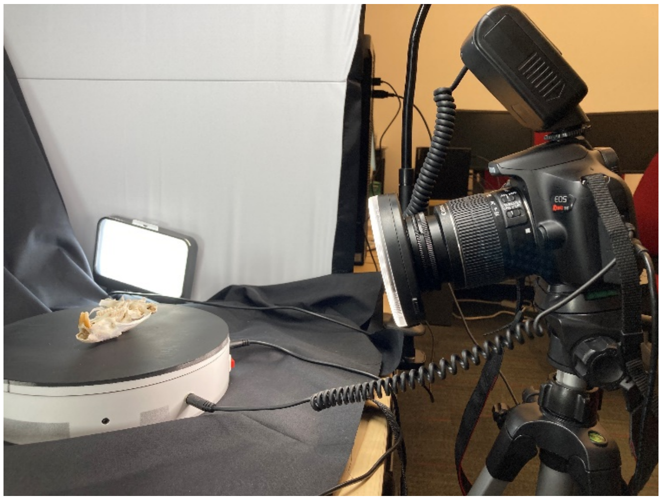

Pictures show the camera set-up in the author CZ's office for this study.

#### 2.2 Camera parameter setup

it is important to keep the entire specimen in focus throughout the photography session. This is best done by manually adjusting the settings on the camera (such as the aperture and focal ring) and creating a depth of field that will keep the specimen in focus in all orientations during photography session. This requires the knowledge of the subject distance to camera, the focal length of the objective and the aperture. These settings can be entered into a depth-of-field (DoF) calculator (e.g., https://www.photopills.com/calculators/dof) to calculate a working distance for the session (see the screenshots below).

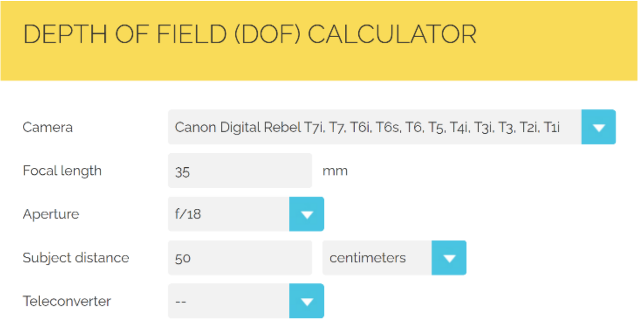


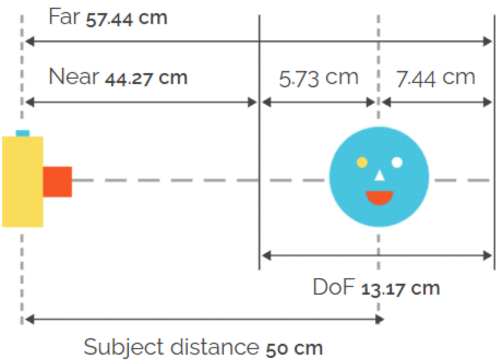

- Subject distance is the distance between the center of the object to the camera sensor, which is marked on the body of the camera (see picture below)

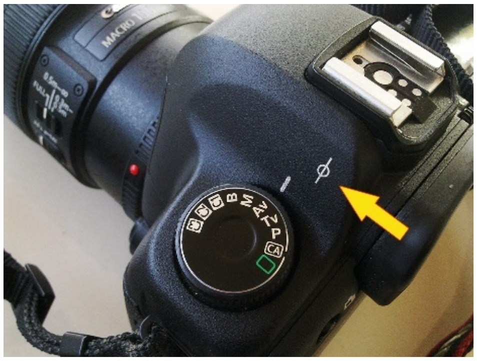

Picture from: https://3.bp.blogspot.com/-tjHpVMrsDeg/UoynJpZzuvI/AAAAAAAAIc4/zmzMtUyCdUc/s1600/focal-plane-mark-on-the-camera-body.jpg

- Ensure the specimen stays within the calculated DoF during the turntable rotation.
- Manually adjust the focus ring to ensure different parts of the specimen are similarly in focus.
- Adjust the shutter speed to control the exposure of photos to an appropriate level.
- We used 12MP setting on our camera (4608 × 3456)
- Once a set of camera parameters is achieved, take a few test photos of the specimen by rotating the turntable to different angles to ensure consistent sharpness of photos.

#### 2.3 Taking photos

For this study, we take six sets of photos. The first five sets are the basis of model reconstruction. The sixth set is optional and is taken with Aruco markers to scale the reconstructed 3D model to its real-life dimensions automatically. Photos of the specimen 39384 are provided as the sample data (https://osf.io/b39yx/, DOI: 10.17605/OSF.IO/B39YX). The folder "39384_raw_photos" contains the raw photos. The folder "39384_masked_photos" contains the masked photos submitted to WebODM (see sections 2.3, 3, and 4). The camera parameters for each set achieved from the DoF calculators are listed in the table below:

| Set | Specimen Orientation | No. of pictures | Focal length (mm) | ISO | Aperture (F stop) | Shutter speed (sec) | Subject distance (in cm) | Depth of field (in cm) |
|---|---|---|---|---|---|---|---|---|
| 1 | Vertical | 48 | 35 | 100 | 13 | 1/6 to 1/8 | 45 | 7.85 (41.41 to 49.27) |
| 2 | Vertical | 48 | 35 | 100 | 13 | 1/6 to 1/8 | 45 | 7.85 (41.41 to 49.27) |
| 3 | Vertical | 48 | 35 | 100 | 13 | 1/6 to 1/8 | 45 | 7.85 (41.41 to 49.27) |
| 4 | Horizontal | 64 | 35 | 100 | 16 | 1/5 | 48 | 10.37 (43.23 to 53.96) |
| 5 | Horizontal | 64 | 35 | 100 | 16 | 1/5 | 48 | 10.37 (43.23 to 53.96) |
| 6 | Horizontal (markers) | 48 | 35 | 100 | 18 | 1/4 | 50 | 13.17 (44.27 to 57.44) |

While they are optimized for our study and specific camera model, they can be used as a starting point for other skeletal elements. Remember to recalculate the DoF every time the specimen to camera distance is changed.

**Set 1.** Place the specimen in a vertical position with the nose facing down. First, fix the nose on the putty on a vertical stand. Adjust the tripod height to make the lens faces approximately the center of the specimen. Take 48 equally spaced pictures (7.5$^{o}$ of rotation at each step).

**Set 2.** Raise the camera to about 12 cm (approximately twice the AP length of the specimen) higher than set 1 and rotate the camera down to approximately face the center of the specimen. Slightly adjust the focal ring if needed. Take 48 pictures.

**Set 3.** Turn the skull upside down and fix the foramen magnum on the putty on the vertical stand. The camera paraments can stay the same. Slightly adjusting the focus ring is needed. Take 48 pictures.

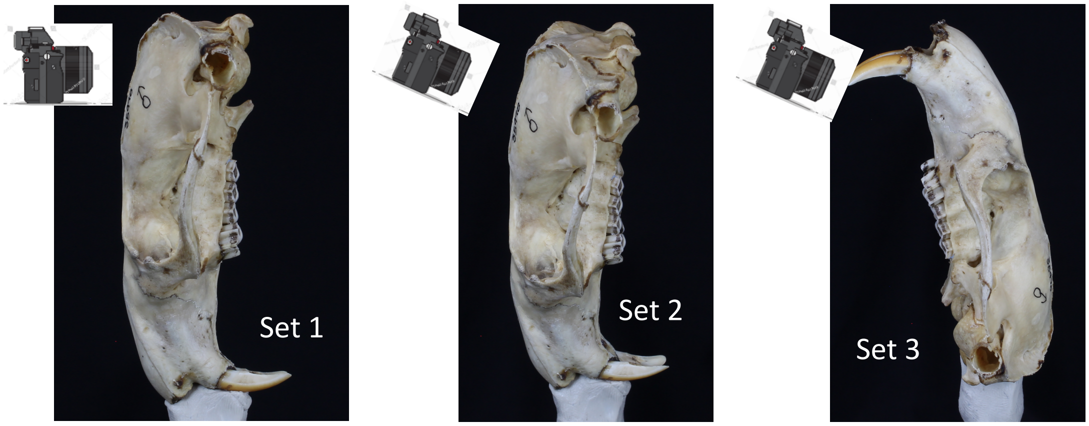


**Set 4.** Put the skull horizontally on the turntable with the ventral surface of the skull facing down and adjust the tripod to point horizontally to the center of the beaver skull. A small platform might be useful to elevate the specimen off the turntable a little. Slightly adjust the focal ring. Take 64 pictures.

**Set 5.** Turn the skull to over to have the ventral surface facing upwards. Take 64 pictures.

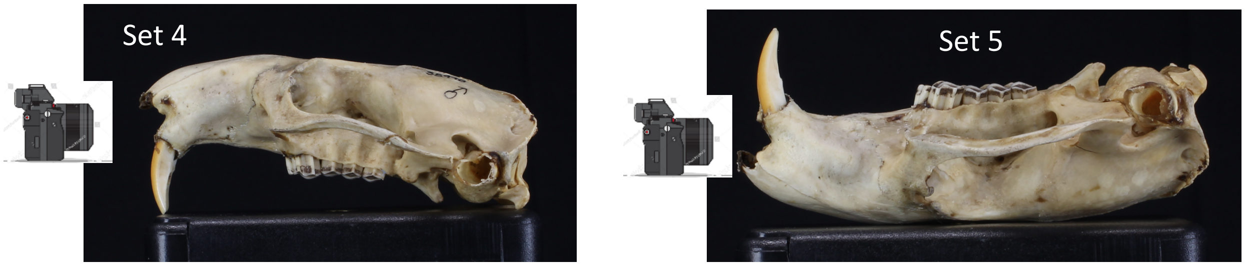


**Set 6.** This set of photos is used for automatedly scaling using the Aruco markers.

- Design a sheet of Aruco markers (e.g., in Microsoft PowerPoint) with markers from ID 0 to ID 9 (markers can be generated at https://chev.me/arucogen/). We created each marker as 8mm wide, which was sufficiently large for our imaging protocol. Put a slightly larger white square below to create a contrast with the black background. An example PowerPoint is provided as part of the SOM.

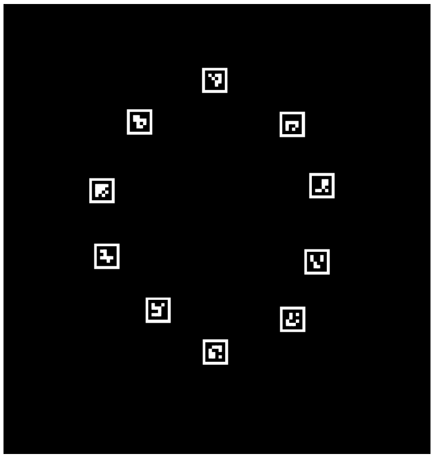

A sample Aruco marker sheet

- Make sure the measurement units are metric, as we will use millimeters to describe the location of each marker in the next step.
- Record the horizontal and vertical position of the upper left corner of each marker from PowerPoint (Size and Position →Position) as the x and y coordinate into a text file (e.g., gcp_coord.txt). This assures that the distances calculated from these coordinates match the actual measurable distances between the markers. Enter these into a text file (e.g., “gcp_coord.txt”) following the convention below:
- The first column is the marker ID. The next three columns are x, y, and z coordinates of the marker (in millimeter units). Note that the z coordinate for each marker is fixed (1.00). This is because the markers are printed on a 2D surface.

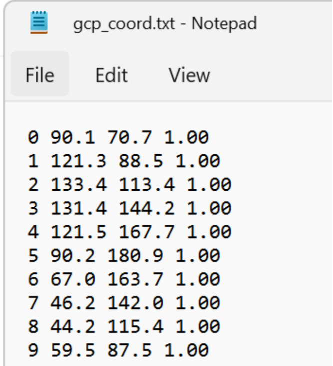

gcp_coord.txt - Notepad

File Edit View

```
0 90.1 70.7 1.00
1 121.3 88.5 1.00
2 133.4 113.4 1.00
3 131.4 144.2 1.00
4 121.5 167.7 1.00
5 90.2 180.9 1.00
6 67.0 163.7 1.00
7 46.2 142.0 1.00
8 44.2 115.4 1.00
9 59.5 87.5 1.00
```

- In the sample data available at https://osf.io/b39yx/ (DOI: 10.17605/OSF.IO/B39YX), the sample “gcp_coord.txt” can be found in the folder “39384_raw_photo” that contains the raw photos.
- When processing, WebODM will assume the unit of input coordinates by default as meter, as that's the convention in geospatial coordinate systems. However, this does not influence actual data analysis-based 3D model because MeshLab and 3D Slicer correctly treats these values as millimeters.
- Cut the marker to a circular shape and put it on the turntable. Use tapes to keep the sheet as flat as possible.
- Put the specimen on the center of the sheet with its base facing down. Raise the camera about seven centimeters higher from the position for the photo sets 5 and 6 to take 48 pictures. Ensure four to five markers are visible in each picture.

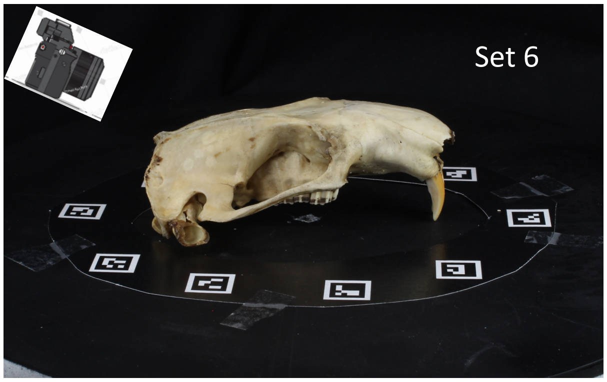


#### 3. Recognize Aruco markers from photos

We use the “Find-GCP” python package (https://github.com/zsiki/Find-GCP) to recognize markers in each photo and record their pixel location of each marker’s upper left corner.

- Install the OpenCV package (https://docs.opencv.org/4.x/df/d65/tutorial_table_of_content_introduction.html) in the terminal through `pip install opencv-python`
- Clone the Find-GCP GitHub repository `git clone https://github.com/zsiki/Find-GCP.git` to the designated directory or directly download the zip file.
- Run the following script in `python`

```
/path/gcp_find.py -t ODM -i /path/gcp_coord.txt --epsg 3857 -o
/path/gcp_list.txt `ls /path/photos_gcp/*.jpg`
```

  - The `/path/gcp_find.py` points to the python script `gcp-find.py` in the cloned `Find-GCP` repository.
  - The `/path/gcp_coord.txt` is the text file that stores the x and y coordinates of the upper left corner of each marker from the marker sheet. See step 6 of section 2.3.
  - EPSG 3857 is a commonly used geographical protocol adopted by ODM.
  - `` `ls /path/photos_gcp/*.jpg` `` lists all photos with markers from the photo (jpg format) set 6 stored in the folder `/path/photos_gcp/`.
  - The `/path/gcp_list.txt` is the output file `gcp_list.txt` stores the recognized marker, and its ID and coordinates in each picture from the photo set 6 of a specimen (see step 6 of section 2.3).
  - A sample output (`gcp_list.txt`) from the script is shown below (this text file can also be found in the sample data in “39384_masked_photos/set6” in the sample data available here: https://osf.io/b39yx/, DOI: 10.17605/OSF.IO/B39YX). The three leftmost columns are the coordinates of the recognized marker from the reference file (gcp_coord.txt) generated in the previous step. The fourth and fifth columns from the left indicate the pixel coordinates of the recognized marker (specifically coordinates of its upper-left corner) in the photo. The sixth column from the left indicates the filename of the photograph marker is recognized from. Finally, the rightmost column is the ID of the recognized marker.

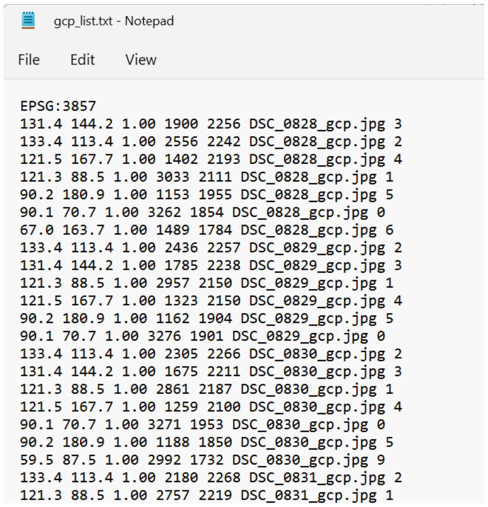

gcp_list.txt - Notepad

File Edit View

```
EPSG:3857
131.4 144.2 1.00 1900 2256 DSC_0828_gcp.jpg 3
133.4 113.4 1.00 2556 2242 DSC_0828_gcp.jpg 2
121.5 167.7 1.00 1402 2193 DSC_0828_gcp.jpg 4
121.3 88.5 1.00 3033 2111 DSC_0828_gcp.jpg 1
90.2 180.9 1.00 1153 1955 DSC_0828_gcp.jpg 5
90.1 70.7 1.00 3262 1854 DSC_0828_gcp.jpg 0
67.0 163.7 1.00 1489 1784 DSC_0828_gcp.jpg 6
133.4 113.4 1.00 2436 2257 DSC_0829_gcp.jpg 2
131.4 144.2 1.00 1785 2238 DSC_0829_gcp.jpg 3
121.3 88.5 1.00 2957 2150 DSC_0829_gcp.jpg 1
121.5 167.7 1.00 1323 2150 DSC_0829_gcp.jpg 4
90.2 180.9 1.00 1162 1904 DSC_0829_gcp.jpg 5
90.1 70.7 1.00 3276 1901 DSC_0829_gcp.jpg 0
133.4 113.4 1.00 2305 2266 DSC_0830_gcp.jpg 2
131.4 144.2 1.00 1675 2211 DSC_0830_gcp.jpg 3
121.3 88.5 1.00 2861 2187 DSC_0830_gcp.jpg 1
121.5 167.7 1.00 1259 2100 DSC_0830_gcp.jpg 4
90.1 70.7 1.00 3271 1953 DSC_0830_gcp.jpg 0
90.2 180.9 1.00 1188 1850 DSC_0830_gcp.jpg 5
59.5 87.5 1.00 2992 1732 DSC_0830_gcp.jpg 9
133.4 113.4 1.00 2180 2268 DSC_0831_gcp.jpg 2
121.3 88.5 1.00 2757 2219 DSC_0831_gcp.jpg 1
```

### 4. Photo pre-processing

#### 4.1 Masking the background

To speed up the reconstruction process and to make it more accurate, it is advisable to remove the background and make the specimen the most prominent structure in the image. Below we describe how to accomplish this using out custom developer scripts in python console of Slicer and its SlicerMorph extension. If the user prefers, same procedure can be done in other image processing programs such as ImageJ or GIMP or many others. However, in that case it is important to make sure that the image dimensions before and after the processing remained unchanged, and that only the background is masked as solid black.

In future these steps will be fully automated in Slicer as a separate extension that we are developing.

Use a custom script (https://github.com/chz31/PhotoGram/blob/main/mask_image_ROI.py) and the ROI tool in 3D Slicer to mask as much background as possible manually. To retain the original dimensions of each picture, color within the masked region is uniformly set to black (0).

- Make sure that the SlicerMorph extension is installed as described in the Prerequisites section. Please make sure your Slicer scene is clear before you start this part.
- Use the ImageStack module from SlicerMorph to import one photo set as a vector volume. Check the “full resolution” option and uncheck “Grayscale” button (see https://github.com/SlicerMorph/Tutorials/tree/main/ImageStacks for the tutorial).
- Use the “Add ROI” button (highlighted in red in the picture below) or go to the “Markup” module to create an ROI (for tutorials, see https://github.com/SlicerMorph/Tutorials/tree/main/Markups_1).
  - Use one ROI for one set of photos. Use the colored handler (colored dots) to adjust the bounding box (ROI). Scroll across the entire imported image stack to make sure that the same bounding box fit to object in each image.
  - Small parts of the specimen can be excluded if shown in other photo sets (e.g., the tip of nose). For the photos of vertically mounted specimens, exclude the putty in each picture is necessary.

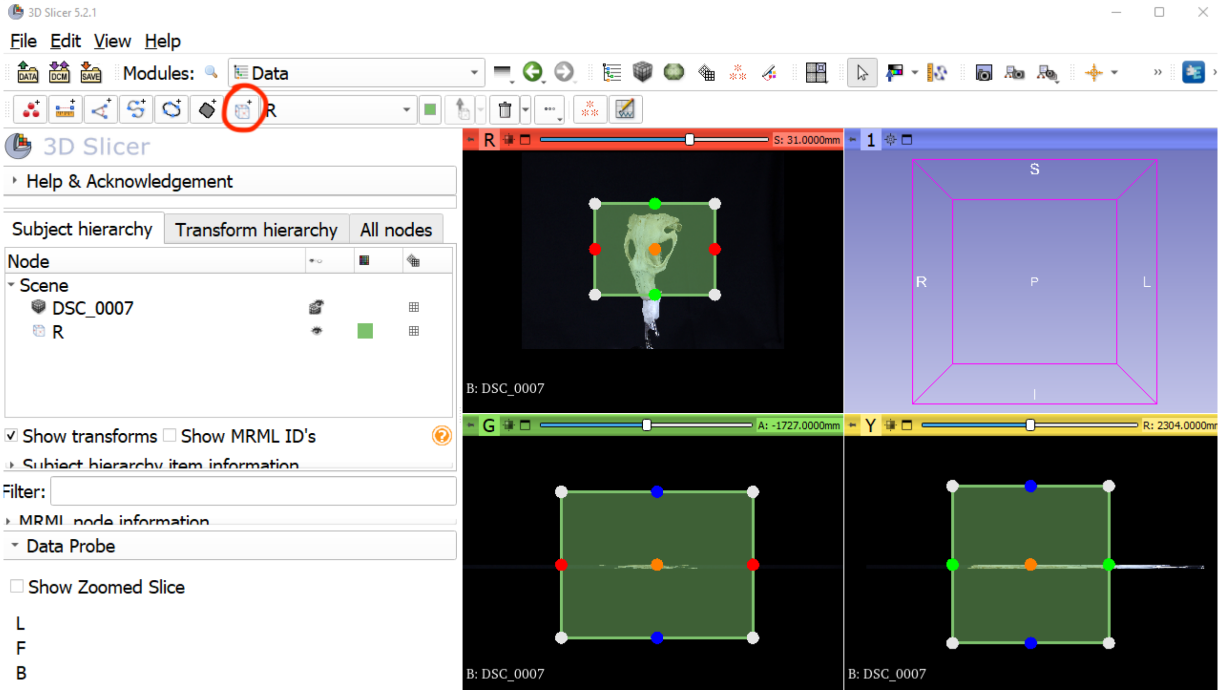

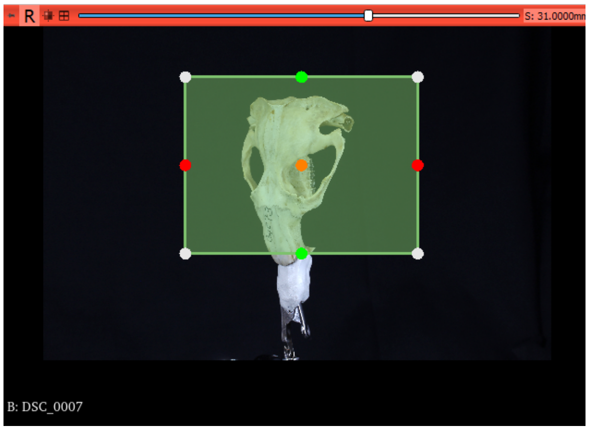

- The photos from the sixth set of photos with Aruco markers (see section 2.3) can also be masked in the same way after the location of each marker is extracted from section 3. Markers should be masked as much as possible or else it may create noise for image registration and model reconstruction.
- Download the custom script [https://github.com/chz31/PhotoGram/blob/main/mask_image_ROI.py](https://github.com/chz31/PhotoGram/blob/main/mask_image_ROI.py).
    - In line 3, enter the path to the downloaded “mask_image_ROI.py” as the “filePath”
    - In line 5, enter the directory to the folder containing one photo set as the “rootPath".
    - In line 6, enter the folder name that contains a particular photo set under the “rootPath” as the “setName”
    - In line 7, enter the volumetric node name after importing a photo set into Slicer using the ImageStacks module.
    - In line 8, enter the ROI name used for masking. By default, the name is “R”.

```
1  """
2  import os
3  filePath = "directory/to/mask_image_ROI.py"
4  #
5  rootPath = "directory/to/the/folder/for/all/photos/of/a/specimen" #the full directory that contains all photos of a specimen
6  setName = "vertical_1" #the folder name that contains a particular photo set under the "rootPath"
7  volNodeName = 'photo_volumetric_node_name' #the imported volumetric node name
8  ROI_name = "Crop Volume ROI"
9  
10 inputPath = os.path.join(rootPath, setName)
11 maskedDir = setName + "_masked"
12 outputPath_masked = os.path.join(rootPath, maskedDir)
13 
14 if not os.path.exists(outputPath_masked):
15   os.makedirs(outputPath_masked)
16 
17 exec(open(filePath).read())
18 """
```

- Copy and paste lines 2 to 17 into the python console of 3D Slicer to run the script.
- The output masked images will be stored in a new folder under the specified “rootPath” as the “setName” plus a postscript “_masked”.
- The sample masked photos from the six sets are shown below.

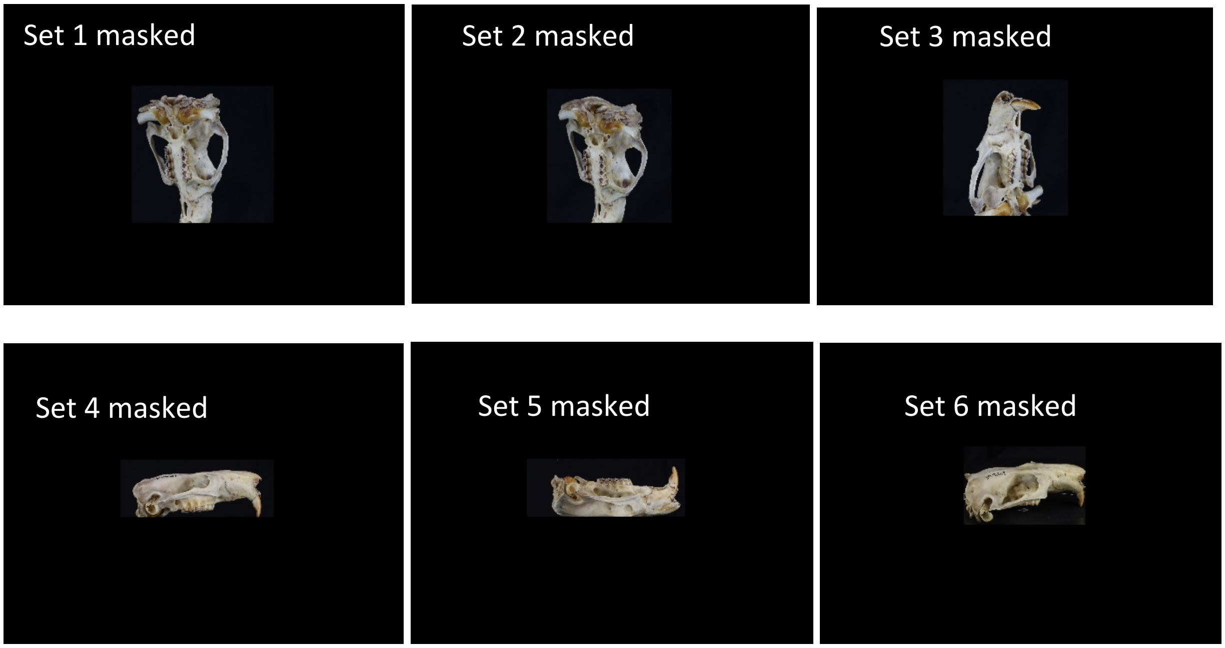


#### 4.2 Use the ExifTool to pass the metadata to the masked photos

Using the ExifTools to pass the metadata from the original photos to the masked photos. Follow the instructions to install ExifTool: https://exiftool.org/install.html. The processed photos from section 4.1 will lost the original metadata after they are saved. ODM requires the camera metadata of photos for successful model reconstruction.

- To transfer the metadata from the original photos to the masked photos from section 4.1, the masked images need to be identically named as the original ones. If using the custom script from section 4.1, the same file names in the output can be guaranteed.
- Run this line of command in the terminal to transfer the metadata:

```
exiftool -tagsFromFile original_photo_dir/%f.%e -all:all
masked_photo_dir/
```

- The `original_photo_dir` is the directory to the folder that contains the original photos with metadata. The `masked_photo_dir` is the directory to the folder that contains the masked images as a result of section 4.1.
- In the sample data available here https://osf.io/b39yx/ (DOI: 10.17605/OSF.IO/B39YX), the masked photos with metadata added can be found in the folder "39384_masked_photos".

### 5. Run 3D model reconstruction and texturing through WebODM

#### 5.1 Submit photos to WebODM

Submit photos to the WebODM interface and run the model reconstruction and texturing.

- Click "Add Project" at the upper right corner of the WebODM web-based interface. Enter project names and descriptions (optional) in the pop-up window and click "Create Project".

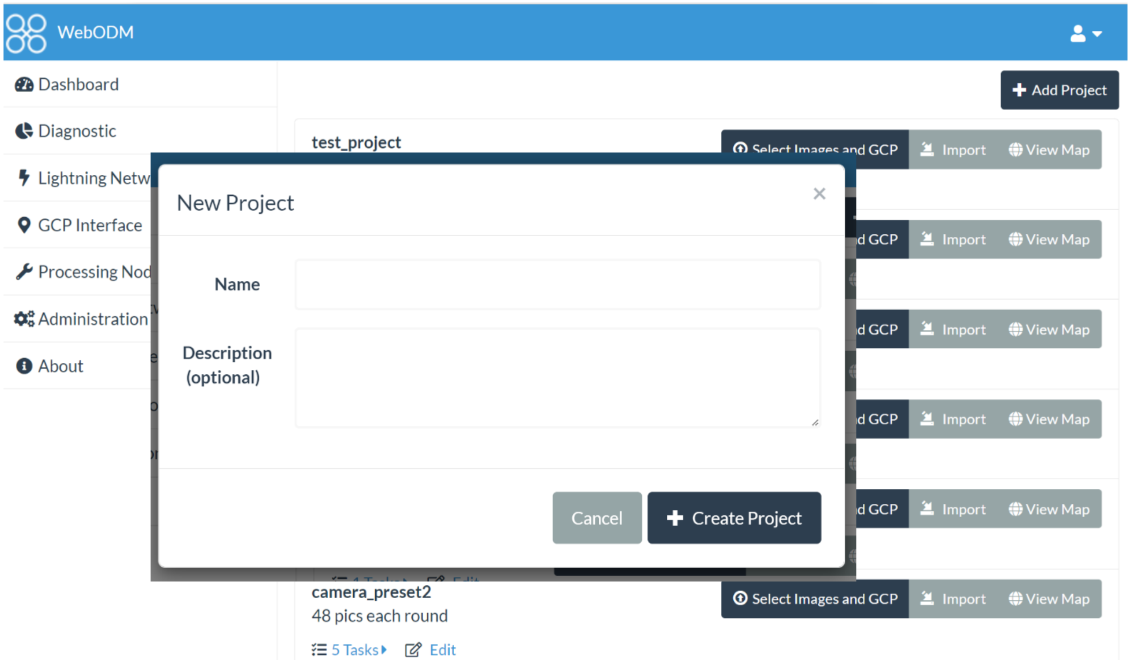


- Click "Select Images and GCP" from the upper right corner in the current project section to upload pictures and initiate a new task. You can change the name of the task. Once a task is added, click "Select Images and GCP" to add more photos from other directories if needed. Remember to select and submit the `gcp_list.txt` generated from section 3 for scaling (the photo set of each specimen should have one unique `gcp_list.txt`). This step is essential to ensure successful output. Click "Edit" in the "Options" entry of the current task after selecting all the photos and the `gcp_list.txt` file to specify parameters. Users can save the parameter settings as a preset and reuse it in other tasks.

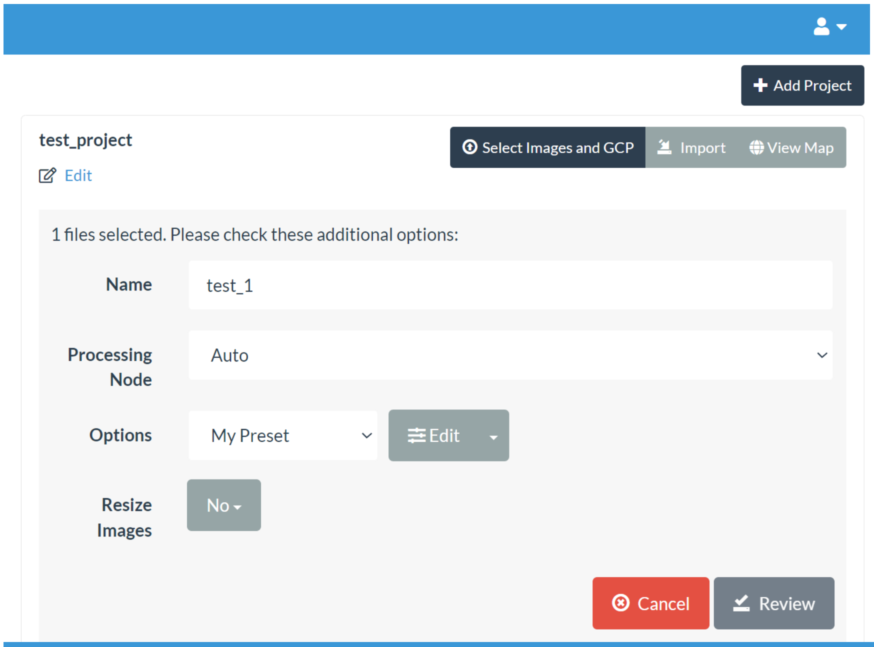

Add Project
test_project
Edit
Select Images and GCP
Import
View Map
1 files selected. Please check these additional options:
Name
test_1
Processing Node
Auto
Options
My Preset
Edit
Resize Images
No
Cancel
Review


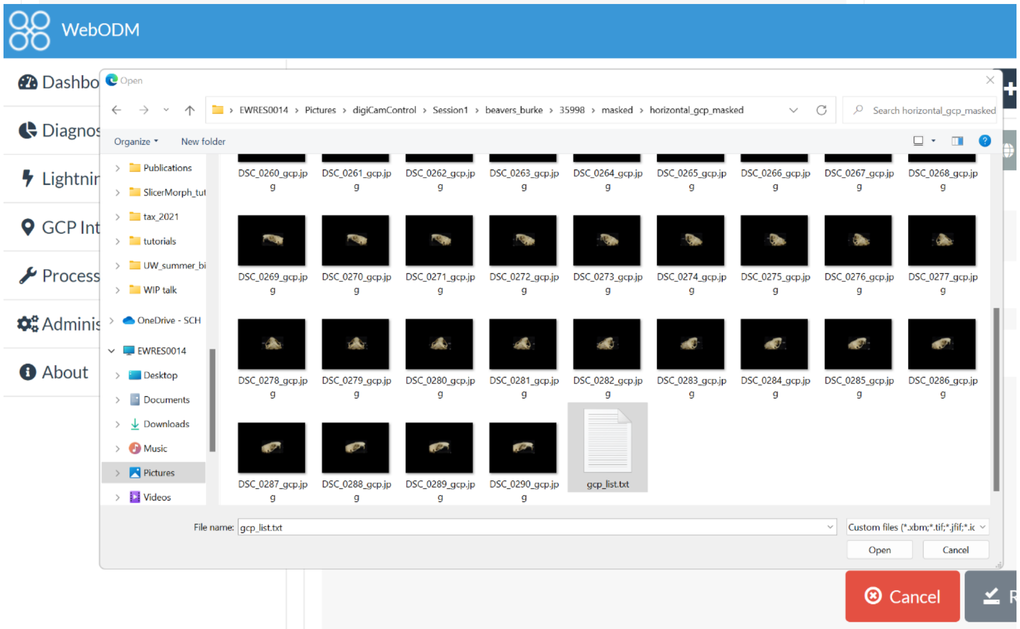

WebODM
Dashbo
Diagnos
Lightnir
GCP Int
Process
Adminis
About
Open
EWRES0014 › Pictures › digiCamControl › Session1 › beavers_burke › 35998 › masked › horizontal_gcp_masked
Search horizontal_gcp_masked
Organize
New folder
Publications
SlicerMorph_tut
tax_2021
tutorials
UW_summer_bi
WIP talk
OneDrive - SCH
EWRES0014
Desktop
Documents
Downloads
Music
Pictures
Videos
gcp_list.txt
File name: gcp_list.txt
Open
Cancel
Cancel


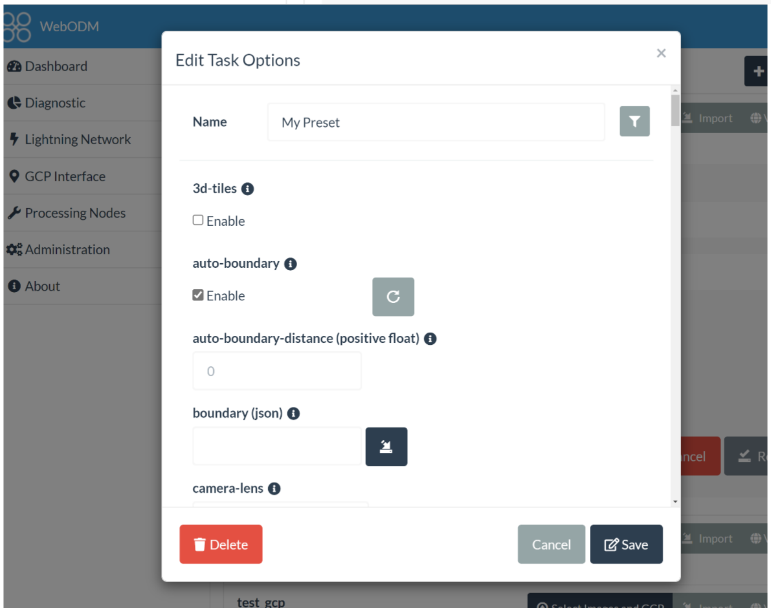

WebODM
Dashboard
Diagnostic
Lightning Network
GCP Interface
Processing Nodes
Administration
About
Edit Task Options
Name
My Preset
3d-tiles
Enable
auto-boundary
Enable
auto-boundary-distance (positive float)
0
boundary (json)
camera-lens
Delete
Cancel
Save
test_gcp

- This is the set of parameters we used for our this study:
    - **auto-boundary: true, bg-removal: true, feature-quality: ultra, mesh-octree-depth: 12, mesh-size: 500000, min-num-features: 30000, no-gpu: true; orthophoto-resolution: 0.3, pc-geometric: true, pc-quality: high, rerun-from: dataset, texture-single-material: true, use-3dmesh: true, verbose: true.**
- Save the changed parameters and press "Review" button to review the parameter set in the "Options" Entry.

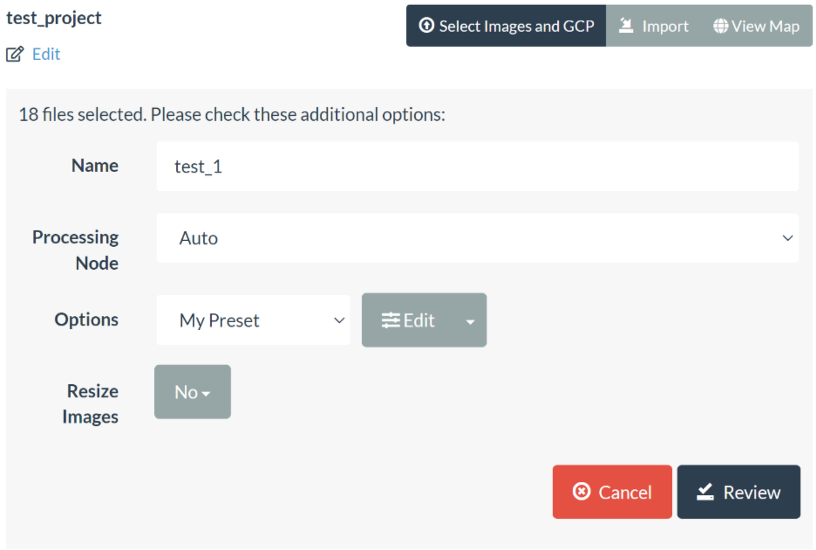


- Click "Start Processing" to start the reconstruction process. Multiple jobs can be submitted to the WebODM queue.

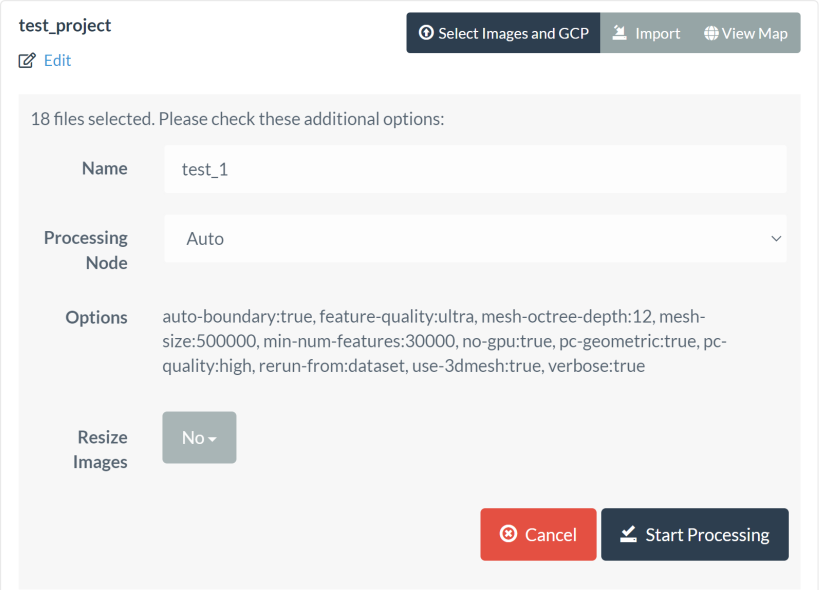


#### 5.2 Parameter setting for WebODM

For detailed explanations of all the parameters, please see the official ODM documentation: https://docs.opendronemap.org/arguments/. We briefly describe the reasonings of our parameter choice below:

- **Bg-removal**: this option is highly recommended to be checked. This option allows ODM to use a deep learning-based algorithm to recognize the object from photos and remove the background. For the current dataset, this option is critical to ensure the consistent success of ODM reconstruction.

- **Texture-single-material**: in order to import the ODM-derived textured model into 3D Slicer for data collection, this option must be checked to produce a single texture image associated with the OBJ model file. Currently, 3D Slicer only supports mapping a single texture image to the model. When unchecked, multiple texture images will be produced.
- **Auto-boundary, pc-geometric, and use-3dmesh**: these three parameters should also be checked as true for biological 3D model reconstruction.
- **Mesh-size**: this parameter determines the maximum vertex count for the output model. We increased the number from the default 200,000 to 500,000 for higher quality models.
- **Feature-quality and min-num-features:** improve feature quality setting and minimum number of features can force the ODM to extract more and higher quality features from images. It may improve image registration for better reconstruction. The feature quality in this study was set as ultra and the min-num-features as 30,000 for better empirically.  Increasing minimum number of features may only increase the computational burden without producing a better outcome. We recommend that users run a few experiments to determine a balance between quality and efficiency.
- **No-GPU:** using GPU can accelerate the process but may lead to lower-quality textured models. Users can experiment with this option for their models. For our study, this option is checked to not use the GPU accelerator.
- **Orthophoto resolution**: this parameter sets up the spatial resolution of the orthophoto and the texture that ODM can produce. The spatial resolution means how large the area a pixel covers (unit: cm/pixel). The default orthophoto resolution is 5 cm/pixel. This generates texture with low resolution. In this study, we input an orthophoto resolution of 0.3 for each task. Here are some suggestions of choosing a proper orthophoto resolution:
    - Overall, the finer the orthophoto resolution (i.e., lower value), the higher the texture's resolution.
    - Note that the orthophoto resolution is capped by an estimated value called ground sampling distance (GSD), an estimated spatial resolution which depends on factors such as camera sensor size, subject distances, camera metadata, and photo resolution. Arbitrarily setting orthophoto resolution to a very low value will not increase the texture and instead result in increased computational burden without any gain.
    - For this study we chose to run WebODM for two to three specimens without inputting the `gcp_list.txt` from section 3. An average estimated GSD for each task can be seen in its drop-down menu. Users also download the full asset and go to the "odm report" folder to see this value in the "odm-report.pdf".

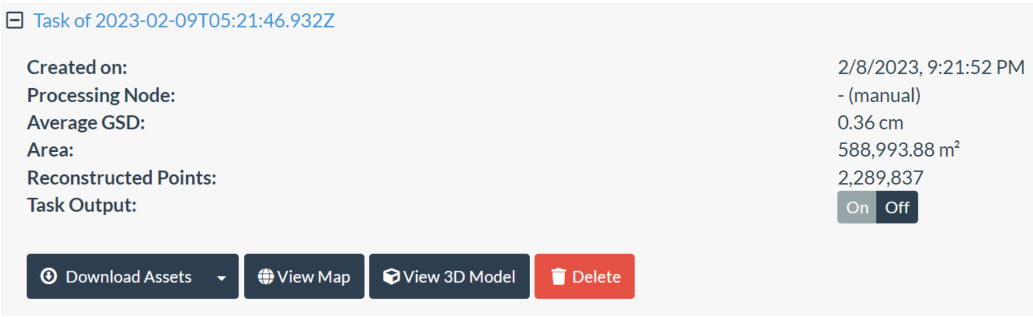

    - **Thus,  we empirically determined that optimum orthophoto resolution for our images was 0.3 using the approach described above. We advise the users to conduct similar experiments with their own data.**
  - For more information about orthophoto resolution, see: https://github.com/OpenDroneMap/ODM/issues/858 and https://community.opendronemap.org/t/lower-resolution-texture-image-after-adding-a-ground-control-point-list-for-scaling/13424.
  - Users can experiment with checking the option **ignore-GSD** if the computational power is high. This will remove the cap of resolution set up by an estimated GSD. It may improve the texture and orthophoto quality in some cases while increasing the computational burden.

#### 5.3 Visualize and download results

- After a task is finished, the output can be visualized in the WebODM interface. Click the “View 3D Model”. It will show the reconstructed dense point cloud.
- Click “Show Model” under the “Textured Model” section at the left control panel to display the textured model. It may take a minute to load the texture map.

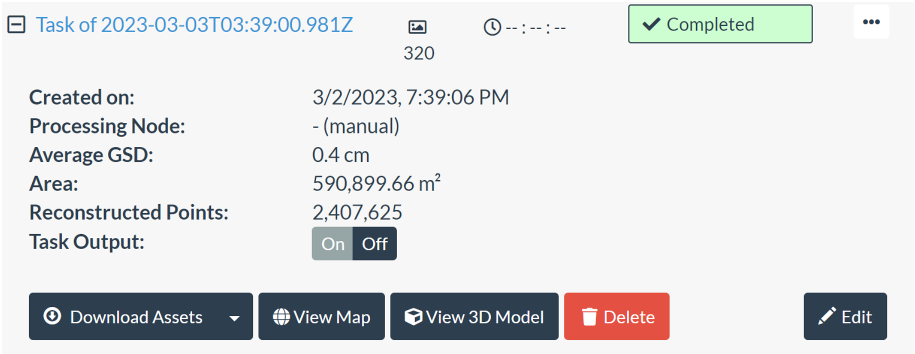


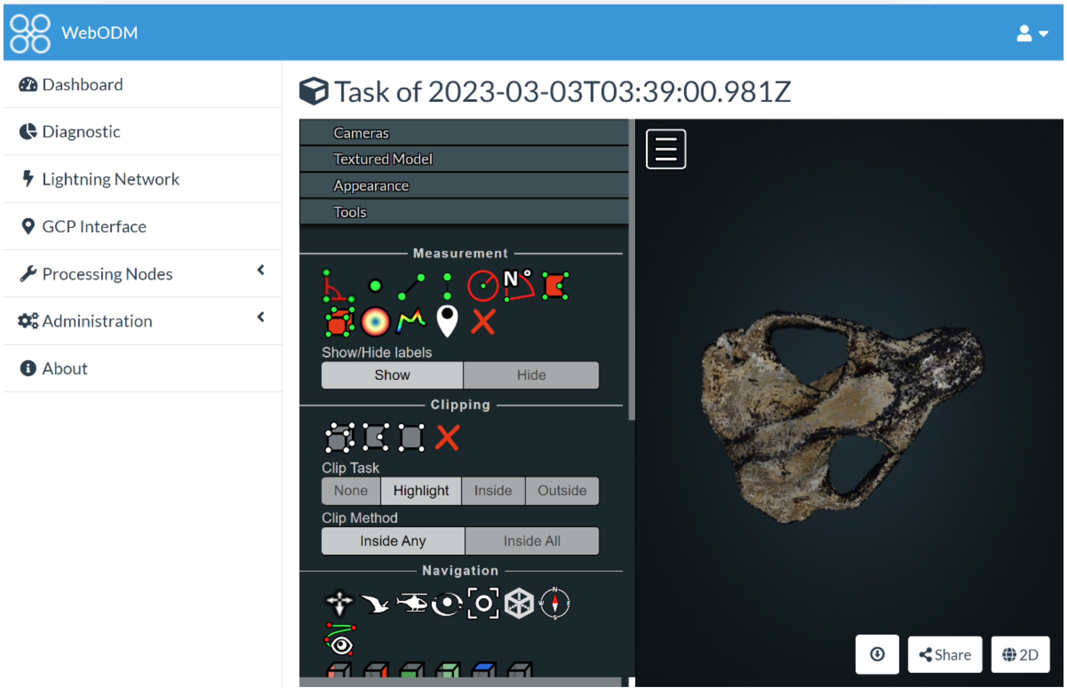

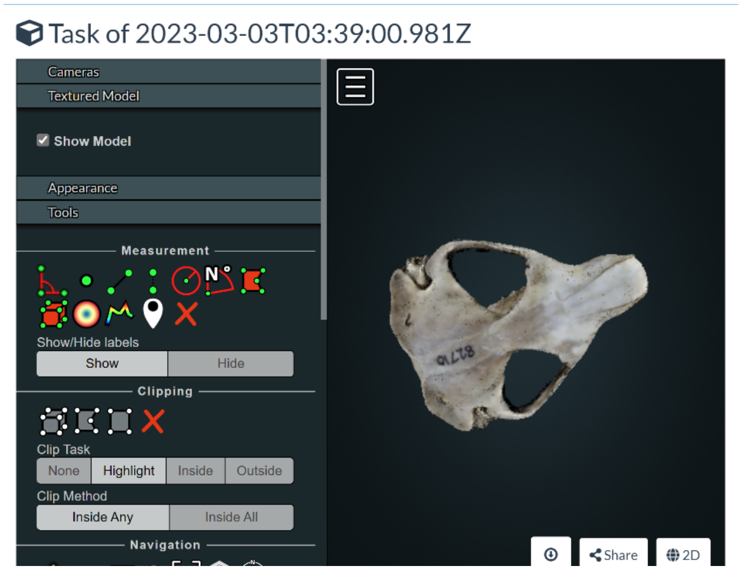


- Check “Show Cameras” under the “Cameras” section on the left control panel to show the reconstructed camera positions. Click each camera position to view the photo.

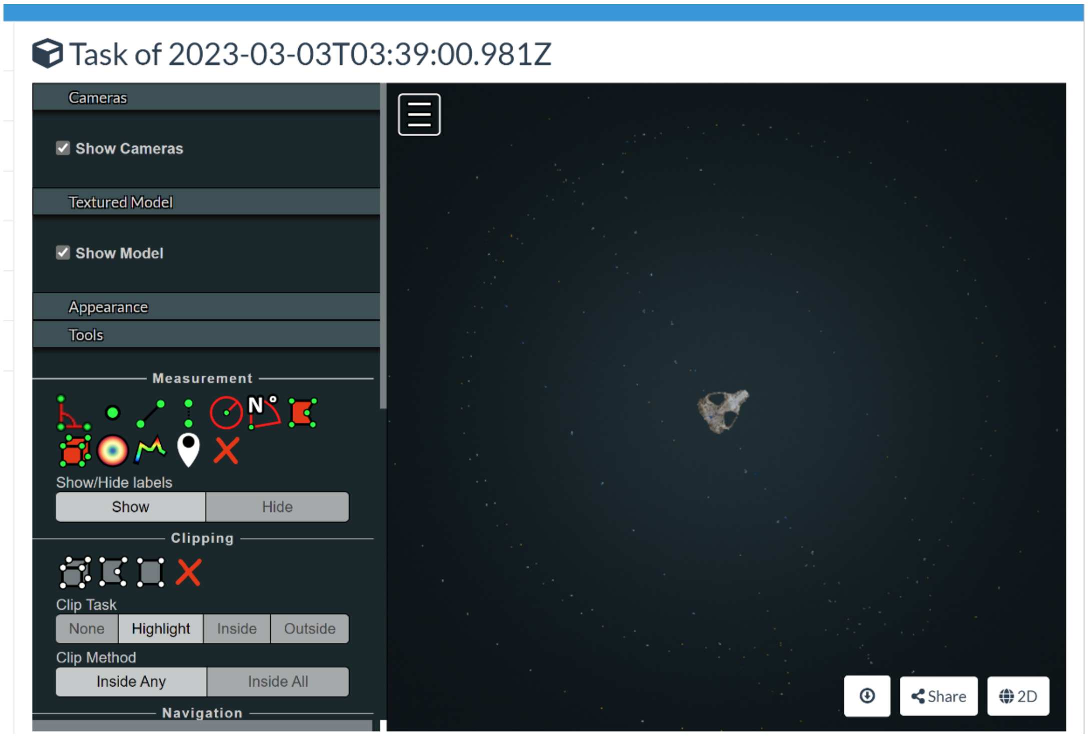


Users can download the textured model or the full asset. Downloading the full asset will download the output from every stage of a WebODM run, as shown in the “Download Assets drop-down menu. For the current study, the “Texture Model” is downloaded as a zip file that contains the 3D model in the OBJ format, an mtl file that stores the texture image information, and the texture image in PNG format.

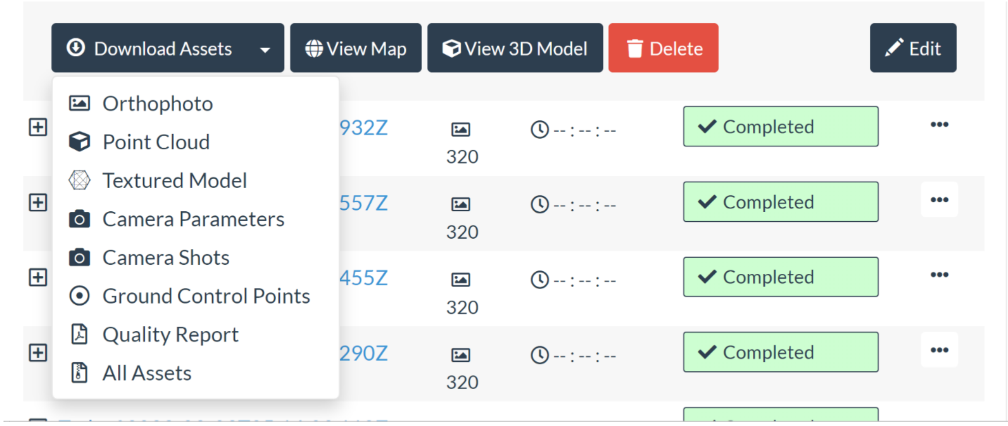


### 6. Model cleaning and data collection

#### 6.1 Model cleaning in MeshLab

Import the OBJ file into MeshLab to clean the textured model.

- Use "Remove isolated pieces (wrt face numbers" (look for "Cleaning and repairing" under "Filters") to clean isolated polygons. The "minimum connected component size" is 1000 for the sample in this study.
- For some polygons floating at the surface (especially the lateral surface of the zygomatic arches like the picture shown below), use "Select vertices" and "Delete selected vertices" from the top toolbar to remove the polygons that connect them with the main component of the model. After that, use "Remove isolated pieces" to remove the isolated polygons.

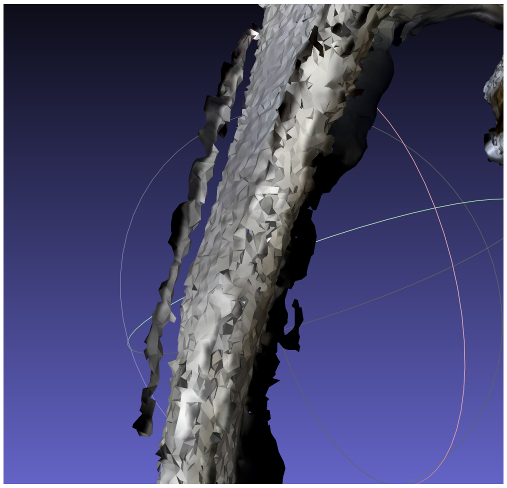

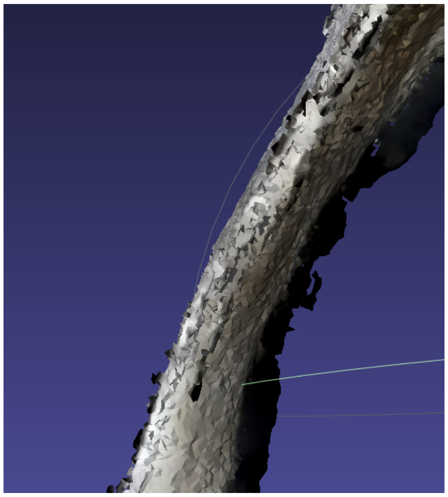

#### 6.2 Import the cleaned textured model into 3D Slicer for data collection.

Please ensure the texture- single-material is checked (see section 5.1 and 5.2) to ensure only one texture map is produced. Currently 3D Slicer only supports mapping one texture image to the OBJ model file. There are two ways of doing this. The first way is to use the Texture Model module in the SlicerIGT extension in 3D Slicer. For a tutorial on the Texture Model module, please see https://github.com/SlicerMorph/Spr_2021/blob/main/Day_1/Models/Models.md.The second way is to import OBJ with automatic texture mapping based on the SlicerMorph extension.

- Ensure that the texture image and the associated mtl file is in the same directory. If downloading the textured model from WebODM, these files will be in the same folder.
- After SlicerMorph is installed in 3D Slicer, drag the OBJ file to 3D Slicer. In the pop-up data dialog, select the “OBJ textured model” option. Click OK will import the OBJ file as a model node in Slicer. The texture image will also be automatically mapped to the model.

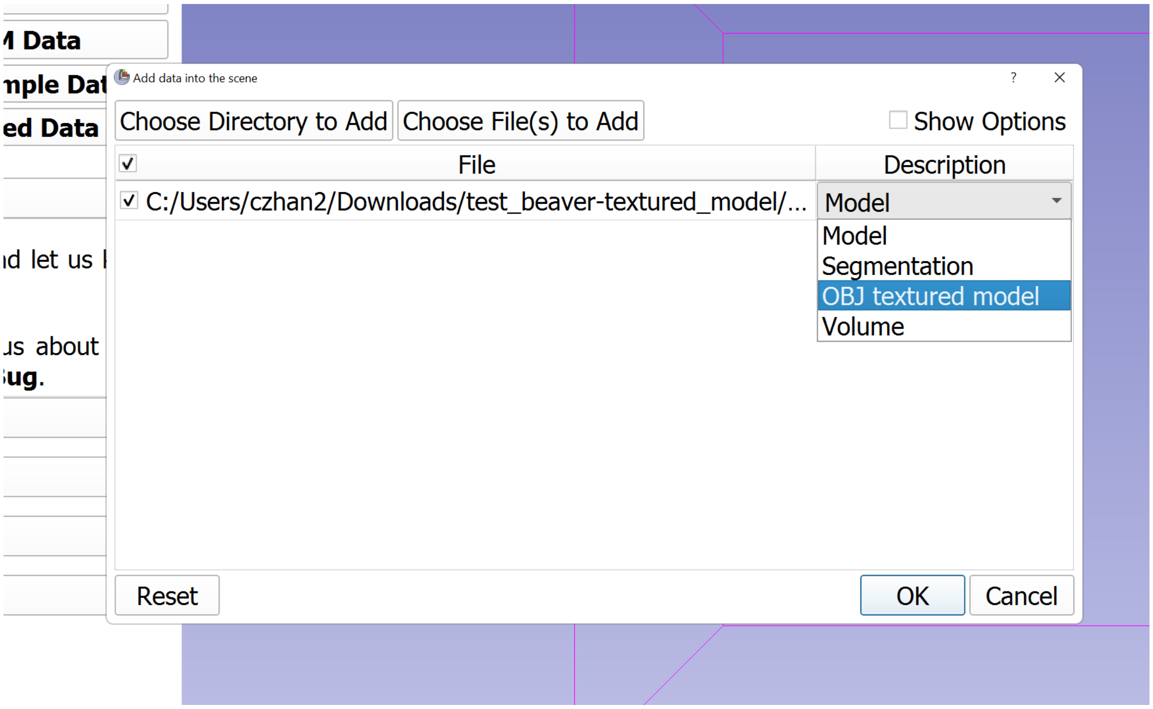
